# Differential analysis of image-based chromatin tracing data with Dory

**DOI:** 10.64898/2026.02.19.706897

**Authors:** Zhaoxia Ma, Miao Liu, Shengyuan Wang, Siyuan Wang, Chongzhi Zang

## Abstract

Spatial organization of the genome plays a vital role in defining cell identity and regulating gene expression. The three-dimensional (3D) genome structure can be measured by sequencing-based techniques such as Hi-C usually on the cell population level or by imaging-based techniques such as chromatin tracing at the single-cell level. Chromatin tracing is a multiplexed DNA fluorescence in situ hybridization (FISH)-based method that can directly map the 3D positions of genomic loci along individual chromosomes at single-molecule resolution. However, few computational tools are available for statistical differential analysis of chromatin tracing data, which are inherently high-dimensional, highly variable and contain many missing values. Here, we present Dory, a statistical method for identifying differential spatial patterns between two groups of chromatin traces. Dory quantifies pairwise spatial distances among genomic regions in a chromatin trace and applies multi-level statistical tests to detect significant structural differences between the two groups of traces. It produces a differential score matrix highlighting region pairs with significant distance difference. Applying Dory to multiple chromatin tracing datasets, we found that the detected chromatin structural changes were associated with alterations in A/B compartments and promoter-enhancer interactions correlated with differential gene expression. Dory is a robust and user-friendly computational tool for quantitative analysis of imaging-based 3D genome data that enables systematic exploration of chromatin architecture and its roles in gene regulation.

## Introduction

Eukaryotic genomes are packed into higher-order chromatin structures in three-dimensional (3D) space^1,2^. This organization affects genome function at multiple levels and is related to gene regulation^3–7^. The high-order chromatin architectures can be measured by high-throughput sequencing-based technologies, represented by Hi-C^8^. Hi-C measures contact frequencies between any two genomic regions and can be used to infer chromatin structures at different scales, such as chromatin loops, topological association domains (TADs), and chromatin compartments. Sequencing-based techniques require enzymatic DNA digestion and measure chromatin contact events rather than the 3D positions of genomic loci^9^. Signal enrichment also often requires a cell population, losing cellular heterogeneity information. Different from sequencing-based techniques, imaging-based techniques can directly measure the positions of genomic regions in a 3D space at the single-molecule level^10–12^. Using multiplexed DNA fluorescence in situ hybridization (FISH), imaging-based techniques can provide the 3D folding pattern of individual chromosomes within a single nucleus. Such techniques include Chromatin Tracing^10^, and its varieties with different genomic resolutions and different levels of multiplexity and multi-modality, such as Optical Reconstruction of Chromatin Architecture (ORCA)^13^, Hi-M^14^, MINA^15^, DNA MERFISH^11^, and DNAseqFISH+^16,17^. Imaging-based 3D genome profiling has been integrated with other single-cell assays, including RNA MERFISH^15^, antibody-based immunoassays^18^, and CRISPR screens^19^, to generate single-cell multi-omic profiles, and has been widely applied across various cell systems and disease contexts^20^. These advanced techniques provide an unprecedented opportunity to study chromatin structures with specific biological functions.

The 3D chromatin structures are different across cell types/states, development stages or diseases^6^, as they have been implicated to associate with gene regulation programs. Differential chromatin interactions can be detected from Hi-C data using computational tools. For instance, diffHic^21^ uses the statistical framework of edgeR^22^, which was designed to identify differentially expressed genes from RNA-seq, to test the differences in Hi-C data between conditions, while HiC-DC+^23^ uses DESeq2^24^, another RNA-seq differential gene expression analysis method, to perform differential analysis on Hi-C contact maps. Other computational tools include HiCcompare^25^ and multiHiCcompare^26^. For single-cell Hi-C computational tools like scHiCDiff^27^ and SnapHiC-D^28^ have been developed to identify differential chromatin contacts. However, these tools are specifically designed for Hi-C data; they are not directly applicable to imaging-based 3D genome profiling data due to the differences in data structure and distribution: Unlike Hi-C, which relies on sequence read counts (integers) to estimate contact frequencies between genomic regions, image-based chromatin tracing produces the 3D coordinates of probed genomic regions, and can directly generate the distance between regions. The distribution of pairwise spatial distances is fundamentally different from that of contact frequencies derived from Hi-C read counts. Furthermore, sequencing depth normalization in methods like HiCcompare^25^ and multiHiCcompare^26^ only applies to Hi-C data and is irrelevant to chromatin tracing data. Therefore, there is a need for computational tools specifically designed for chromatin tracing data.

Several computational tools have been developed for analyzing chromatin tracing data. For example, jie^29^, a spatial genome aligner, is designed to map the image signals to a reference DNA polymer model and to select the true fluorescence spots corresponding to genomic regions. SnapFISH^30^ is designed to identify chromatin loops from multiplexed DNA FISH data. SnapFISH-IMPUTE^31^ is method for imputing missing loci from such image data. However, there is currently no computational tool specifically designed for statistical differential analysis of chromatin structures for imaging-based 3D genome data.

Three major challenges exist in differential analysis of chromatin tracing data. First, due to technical limitations, there can be a high number of probed loci that are not detected across thousands of chromatin traces, leading to missing values in the data^12,30^. Second, chromatin structures are highly dynamic across individual cells^4^. An effective method should account for the high variances in the 3D structure of thousands of chromatin traces from one dataset. Third, the high dimensional nature of chromatin tracing data due to the high degrees of freedom of 3D chromatin traces demand appropriate statistical approaches for robust evaluation of statistically significant differences.

To overcome these challenges, we introduce Dory (<u>D</u>ifferential chr<u>o</u>matin tracing anal<u>y</u>sis), a computational tool for differential analysis of chromatin tracing data. Dory conducts statistical tests to compare two sets of chromatin traces and robustly identify region pairs with differential distances between the two conditions. By applying Dory to several chromatin tracing datasets generated from various biological samples, we show that changes in chromatin structures identified by Dory are partially linked to alterations in A/B compartment interactions and associated with differential gene expression. We also demonstrate that Dory can help identify chromatin regions carrying functional regulatory elements under specific biological conditions, providing insights into gene regulatory mechanisms.

## Results

### The Dory algorithm

Dory is a computational method designed for differential analysis of chromatin structures between two sets of chromatin traces generated from image-based 3D genome profiling techniques (Fig. 1a). The input data are two sets of chromatin traces. Each chromatin trace consists of the 3D coordinates for the probed genomic regions from the experiment, usually for one chromosome. For each chromatin trace, Dory first calculates the Euclidean distance between every pair of probed genomic regions and generates a pairwise distance matrix. Dory then uses the non-parametric one-sided Wilcoxon rank sum test to compare the Euclidean distances of a region pair across all chromatin traces between the two sets (labeled as Condition 1 and Condition 2), generating two p-values for the “greater” and “less” directions, respectively, for each region pair (every non-diagonal element in the pairwise matrix). Dory then generates a differential score (DiffScore), whose absolute value is defined as the negative log10-transformed smaller p-value between the two, and the sign is assigned positive/negative if the smaller p-value is for the “greater” / “less” direction, respectively. Dory calculates the DiffScore for each region pair in the chromatin trace and generates a DiffScore matrix (Fig. 1b), where a positive score indicates that the spatial distance of a region pair is greater in Condition 2 than in Condition 1, while a negative score indicates the opposite. Region pairs that meet a p-value threshold are identified as differential region pairs (DRPs), for each direction (Fig. 1c). The output of Dory includes a DiffScore matrix and identified DRPs comparing two sets of chromatin traces.

**Figure 1.**
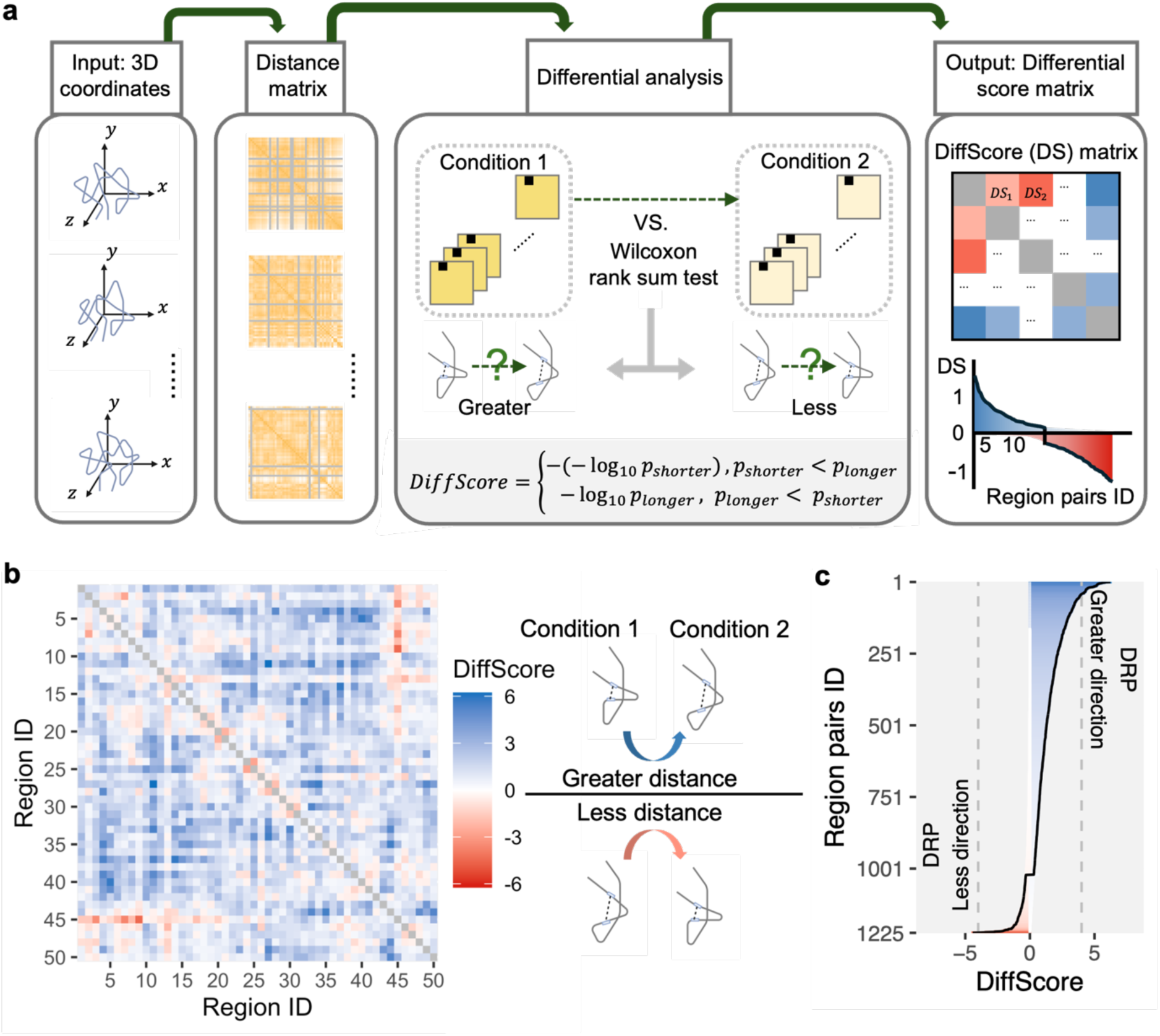
Schematic of Dory. **a**, Schematic representation of the Dory workflow. Dory takes in the 3D coordinates of probed genomic regions from individual chromatin traces as input. In the preprocessing step, each chromatin trace is represented by a distance matrix, where rows and columns represent genomic regions, and each element represents the Euclidean distance between a pair of regions. Missing values are shown in gray. The differential analysis is implemented by applying Wilcoxon rank sum test to the distance vectors of each region pair across all traces between two conditions. The output of Dory includes a DiffScore matrix and identified Differential region pairs (DRPs). **b**, DiffScore matrix. Rows and columns represent probed genomic regions (labeled by IDs), and each element represents the DiffScore comparing Condition 2 versus Condition 1. A positive DiffScore indicates that the distance is greater in Condition 2 than in Condition 1, whereas a negative DiffScore suggests the opposite. The higher the absolute values of DiffScore, the greater difference of chromatin structures between two conditions. **c**, DRPs identified by Dory. DRPs in the “greater” direction represent region pairs with significantly increased distances in Condition 2 than in Condition 1, whereas DRPs in the “less” direction represent region pairs with significantly decreased distances in Condition 2 than in Condition 1.

### Dory exhibits high robustness

To assess the performance of Dory, we used a published dataset of chromatin tracing combined with RNA MERFISH in the mouse fetal liver^15,32^. We used the RNA MERFISH data to determine the cell type for each chromatin trace, and obtained 1,490 and 2,615 chromatin traces for proerythroblast and erythroblast cell types, respectively (Fig. 2a). Each trace consists of the 3D coordinates of 50 genomic regions along Chromosome 19. To investigate the impact of trace count on DiffScores for the 1,225 unique pairs of the 50 regions, we randomly sampled a subset of traces from each cell type. We used Dory to generate the DiffScore matrix between the two cell types using the whole sample and the subsample separately, and calculated the Pearson correlation coefficient (PCC) of the region pairs’ DiffScores between the whole sample and the subsample (Fig. 2b). We subsampled the traces ranging from 100 to 1,000 from the whole sample (Fig. 2b, Supplementary Fig. 1). For each subsample size, we performed 20 independent random samplings to account for variability introduced by random selection. Then, we applied Dory to the two cell types of chromatin traces for each subsample and calculated the PCC of region pairs’ DiffScores between each subsample and the whole sample. The subsamples show consistently high correlations with the whole sample across all 10 cases (i.e., 100, 200, …, 1000 traces) (Fig. 2c). As expected, subsamples with more traces tend to have higher PCC with the whole sample. Notably, the median PCC reaches around 0.7 with as few as 300 sub traces, indicating the robustness of Dory for even small sample sizes (Fig. 2c).

**Figure 2.**
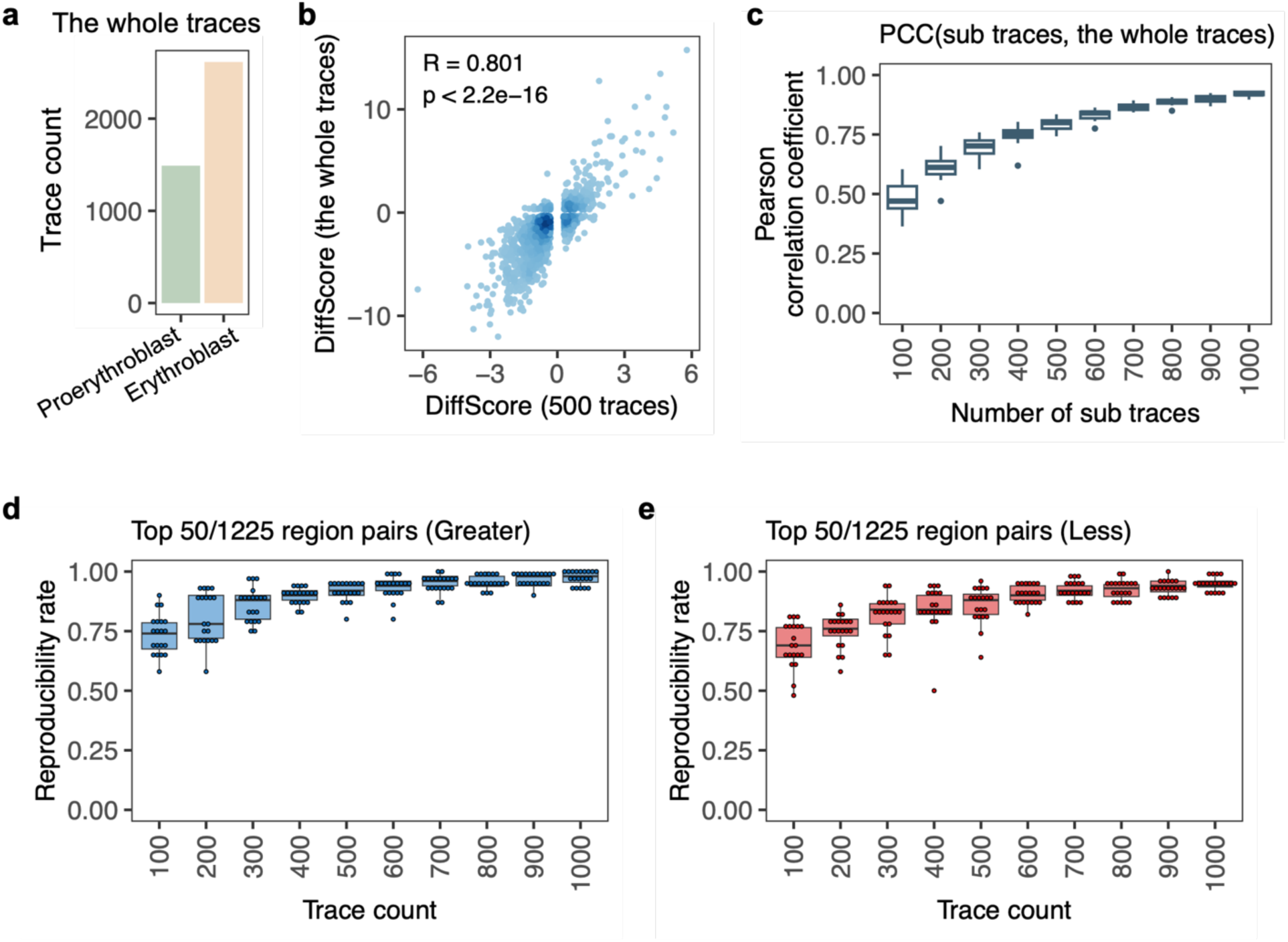
The robustness evaluation of Dory. **a**, Bar plot showing the total trace counts for proerythroblast (1490 traces) and erythroblast (2615 traces) samples. **b**, Scatter plot displaying the correlation of the DiffScore calculated from the whole trace samples versus a subset of 500 traces for each cell type. DiffScores for region pairs are generated by comparing erythroblasts to proerythroblasts. **c**, Box plots of Pearson correlation coefficients (PCC) between DiffScores calculated from the whole sample and the subsampled datasets of varying sizes (100, 200, …, 1,000 traces for each cell type). Each box represents 20 independent random samplings; each data point in the box corresponds to the PCC between the whole sample and one random sampling. **d, e**, Reproducibility rates for the top 50 of 1,225 region pairs with greater (**d**) or less (**e**) genomic distances in erythroblasts compared to proerythroblasts, evaluated across different trace counts. See Methods for details on the calculation of reproducibility rates.

To evaluate the robustness of Dory in reproducibly identifying the most significant DRPs, we defined a reproducibility rate as the proportion of the top 50 DRPs from one subsample that also occur among the top 50 DRPs from any of the other 19 subsamples in the same direction. We calculated the reproducibility rate for each subsample for both directions: DRPs whose distance is greater in erythroblast than in proerythroblast (Fig. 2d) and those whose distance is less in erythroblast than in proerythroblast (Fig. 2e). As a result, approximately at least 70% of the DRPs are reproducible even at the smallest subsample with only 100 traces. As expected, subsamples with a higher number of traces have a higher reproducibility rate.

Taken together, these results show that while the increase of trace count may lead to increase in Dory’s performance, Dory can reproducibly identify the most significant DRPs even with a few hundred chromatin traces. Dory is a robust method that is effective and reproducible for DRP identification.

### Dory identifies large-scale chromatin architectural changes that are partially associated with alterations in A/B compartment organization

To interpret the Dory results and to understand the biology implied by Dory analysis, we used the same chromatin tracing dataset from mouse fetal liver^15,32^, which contains six cell types annotated based on RNA MERFISH, to investigate the differential chromatin structures across cell types. The 50 probed genomic regions along Chromosome 19 have an approximately 1Mb resolution, corresponding to 50 TADs across Chromosome 19. We used Dory to compare the chromatin traces between every two cell types, generating DiffScore matrices. Additionally, we examined the A/B compartment structures in each cell type from the chromatin tracing data using the same computational workflow for calculating A/B compartment scores from Hi-C data^10^, and calculated the A/B interaction score for each region pair by multiplying the compartment scores. A pair of regions belonging to the same type of compartment will have a positive interaction score, while a pair of regions belonging to different types of compartments will produce a negative interaction score. Then we calculated the difference of A/B interaction scores between every two cell types for each region pair, denoted as ABchanges (Fig. 3a, see details in Methods). A region pair with a positive ABchange with a large absolute value tends to shift into the same compartment (e.g., AB to AA, and AB to BB), whereas a region pair with a negative ABchange with a large absolute value tends to shift into different compartments (e.g., AA to AB, and BB to AB) (Supplementary Fig. 2a, b). We observed a negative correlation between Dory-calculated DiffScore and ABchange comparing every two cell types (Fig. 3b, Supplementary Fig. 3), with high significance demonstrated by permutation test (Fig. 3c, Supplementary Fig. 4). This significant correlation indicates that most Dory-identified DRPs are associated with chromatin compartment interaction changes: a negative DiffScore reflects decreased spatial distance between regions, corresponding to increased A/B interactions and a tendency for the interacting regions to reside in the same compartment, and vice versa (Fig. 3b).

**Figure 3.**
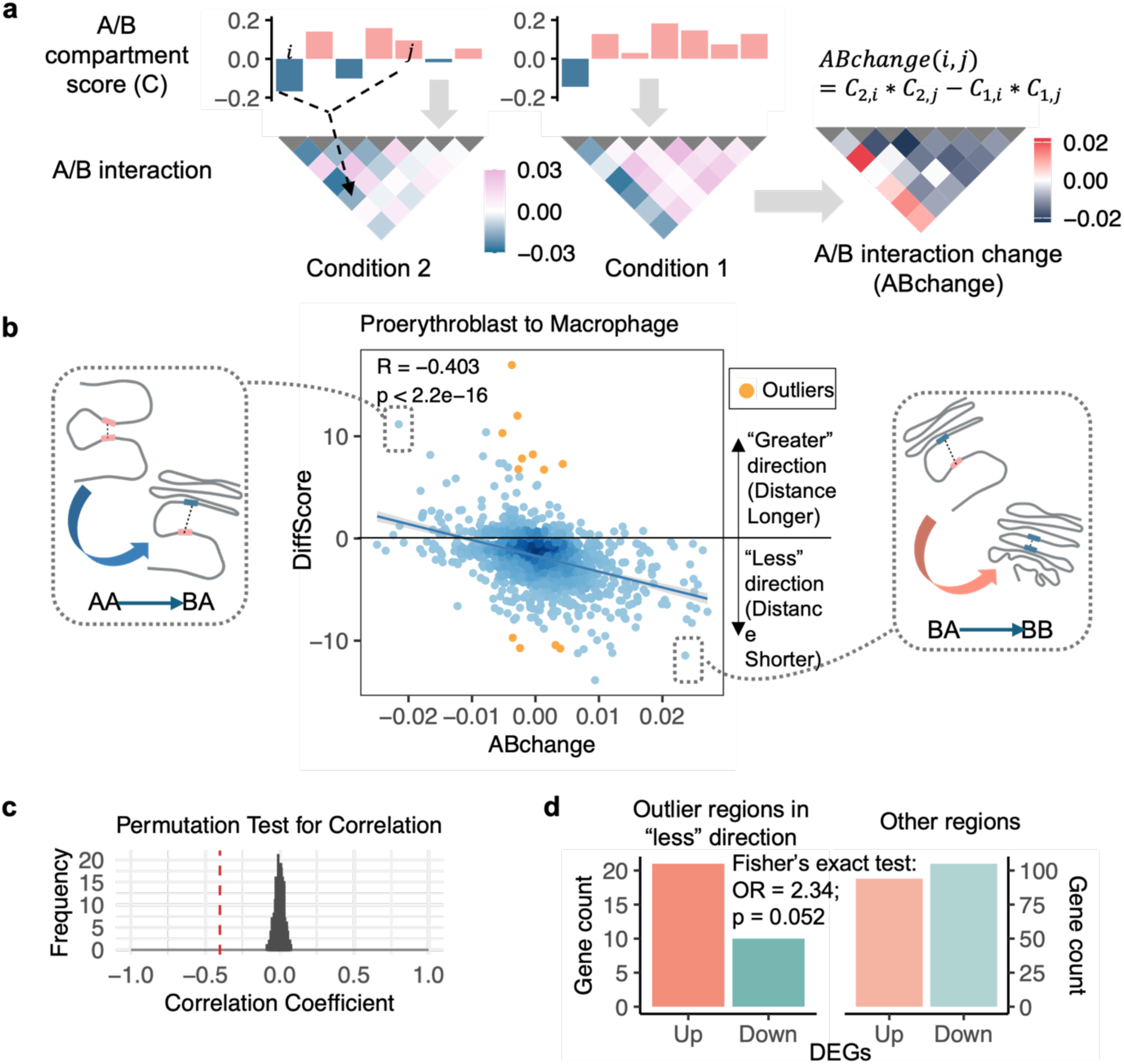
Chromatin structure changes are partially associated with alterations in A/B compartment interactions. **a**, Schematic of A/B interaction change score (ABchange). The upper panel represents the A/B compartment scores of regions in conditions 1 and 2. The lower left panel represents each region pair’s A/B interaction score, calculated as the product of the compartment scores for the two regions in each condition. The lower right panel represents the ABchange, defined as the difference in A/B interaction scores between the two conditions. **b**, Scatter plot displaying the correlation between the DiffScore and ABchange for the 1225 unique pairs from the 50 regions along Chromosome 19, comparing macrophages with proerythroblasts. The increase in A/B interaction is associated with the decrease in the spatial distance between region pairs. Two representative data points (dashed boxes) illustrate this trend: region pairs moving to closer tend to shift into the same compartment and exhibit a high positive ABchange (right); region pairs moving to farther tend to shift into different compartments and exhibit a negative ABchange(left). The outliers are highlighted in orange. **c**, Permutation test for the correlation between DiffScore and ABchange. The null distribution (around 0) is the correlations from 1,000 samples of randomly permuted ABchanges. The red dashed line indicates the observed correlation coefficient between DiffScore and ABchange (in **b**). **d**, Numbers of upregulated and downregulated differentially expressed genes (DEGs) in the outlier regions in the “less” direction (left) and in the other regions (right). P-value was calculated by Fisher’s exact test.

While a significant correlation between DiffScore and ABchange across region pairs was observed, this correlation is not perfect. We then further examined the outliers in their relationship (Orange dots in Fig. 3b, see details in Methods). Although distance changes of these outlier region pairs are not associated with their A/B interaction changes, those regions with shorter distances to its paired regions were more enriched for up-regulated genes than down-regulated genes (Fig. 3d), indicating potential ultra-long-range enhancer-promoter interaction and transcription activation^33^, which may happen in active chromatin regions without substantial changes in compartmental structures. These results suggest that the chromatin structure changes at the mega base scale identified by Dory are largely associated with changes in chromatin compartment interactions, while individual loci may also associate with long-range cis-regulatory interactions independent of the compartment organization.

### Dory identifies active enhancers bound by cell type-specific transcriptional regulators

We then sought to demonstrate Dory’s ability to study fine-scale promoter-enhancer interactions on high-resolution chromatin tracing data. *Scd2* is a crucial gene for lipid metabolism during early liver development^34^ and is specifically expressed in hepatocytes^15^. We examined the chromatin tracing data that probed 19 genomic regions upstream of *Scd2* locus with a 5kb resolution in hepatocytes and other cell types in mouse fetal liver^15^. There are 1491 chromatin traces for hepatocytes and 4185 traces for other cell types. We labeled the probed genomic regions sequentially from 1 to 19, with Region 1 as the most distal upstream and Region 19 at the *Scd2* promoter. In the original study^15^, Region 16 was reported to include active enhancers that interact with *Scd2* promoter in hepatocytes. To identify other potential enhancers of *Scd2* in hepatocytes, we utilized Dory to identify regions that are significantly closer to *Scd2* promoter (Region 19) in hepatocytes than in other cell types. Under the p-value threshold of 0.05 (i.e., | DiffScore | = -log10(0.05)), we identified five probed regions including Region 16 that are significantly closer to Region 19 in hepatocytes than in other cell types (Fig. 4a), with their distance changes and DiffScores shown in Fig. 4b. Interestingly, 3 of these 5 regions contained several enhancer-like signatures (ELS) based on the ENCODE^35^ annotation (Fig. 4c).

**Figure 4.**
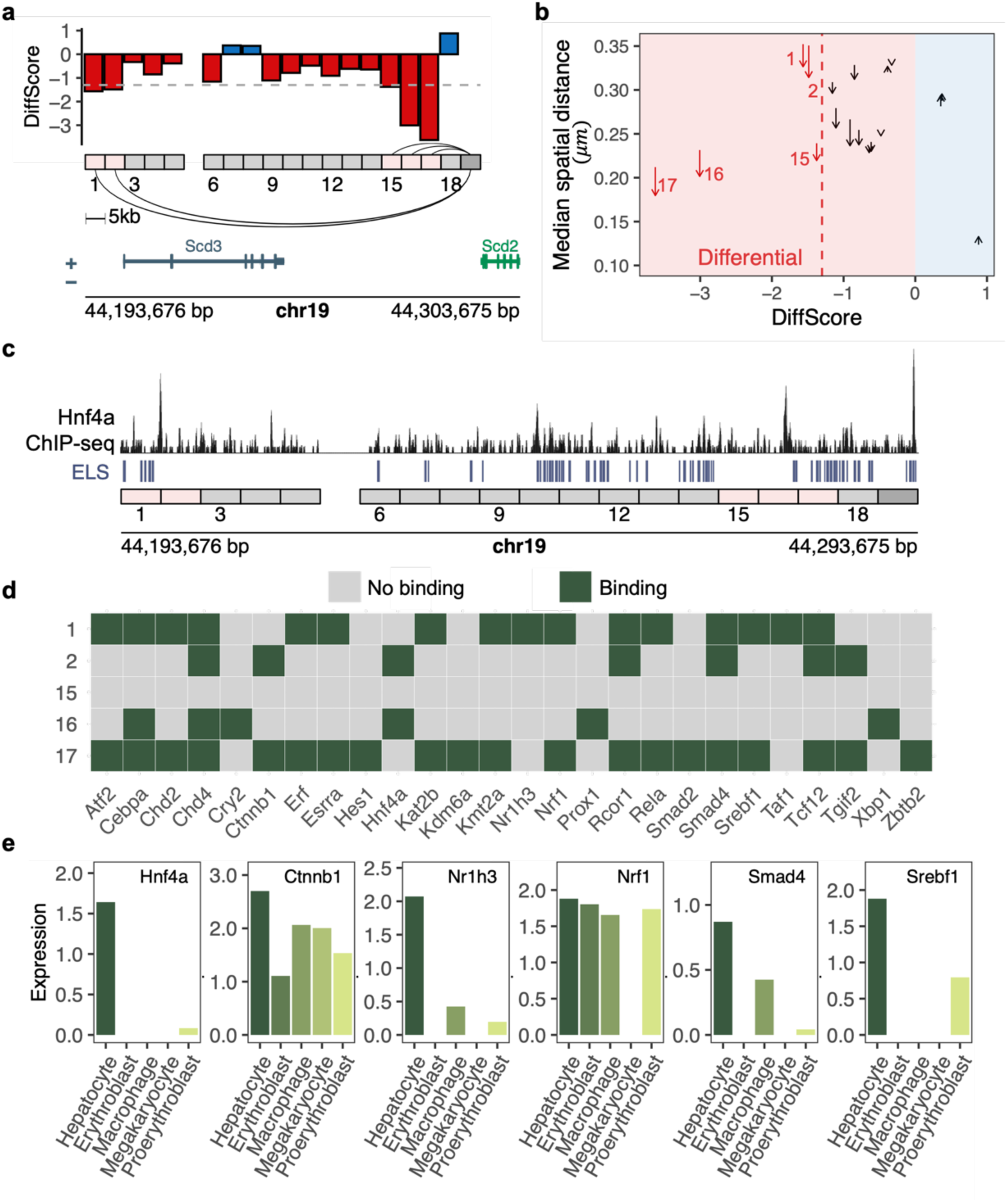
Dory-identified CloserRegions contain potential enhancers of *Scd2* in hepatocytes. **a**, Bar plot displaying DiffScores for region pairs of *Scd2* promoter (Region 19, indicated by dark gray box) and 18 upstream regions in hepatocytes versus other cell types. Regions 1, 2, 15, 16, and 17 are identified as CloserRegions (pink boxes), indicating significantly shorter spatial distances to the *Scd2* promoter in hepatocytes than in other cell types. **b**, Arrow plot displaying distance patterns between *Scd2* promoter and the 18 upstream regions in hepatocytes compared to other cell types. The x-axis represents the DiffScore. The y-axis represents the median distance across all traces. Each region pair is represented as an arrow from non-hepatocyte cell types (arrow start) to hepatocytes (arrow end). The red dashed line indicates the DiffScore threshold. The red arrows are the five CloserRegions. **c**, Hnf4a ChIP-seq signals in mouse liver (GSM2406340), enhancer-like signatures (ELS) from the ENCODE project, and the Dory-identified CloserRegions (pink) among the probed regions near the *Scd*2 locus. The *Scd2* promoter region is marked in dark gray. **d**, Regulator (x-axis) binding sites on CloserRegions (y-axis). Green boxes indicate binding; gray boxes indicate no binding. **e**, Normalized expression levels of identified potential regulators of *Scd2* in hepatocytes and 4 other cell types in mouse fetal liver from the Mouse Cell Atlas.

To identify the potential transcriptional regulators (TRs, transcription factors and chromatin regulators) of *Scd2* that bind in these five regions significantly closer to Scd2 promoter in hepatocytes (hereafter referred to as “CloserRegions”), we utilized TR ChIP-seq data collected in the BART library^36,37^ and their expression profiles from the Mouse Cell Atlas^38^. The BART library contains 5851 ChIP-seq profiles for the binding patterns of 565 unique mouse TRs. We identified 26 TRs that potentially bind in the CloserRegions (Fig. 4d) and exhibit the highest expression in hepatocytes among the five cell types (Supplementary Fig. 5) as potential regulators of *Scd2* in hepatocytes (See details in Methods). For example, the ChIP-seq profile of Hnf4a in liver showed that Hnf4a binds in Regions 2 and 16 (Fig. 4c). Hnf4a is specifically expressed in hepatocytes but not in the other four cell types (Fig. 4e), indicating Hnf4a is a TR specifically active in hepatocytes and potentially regulates *Scd2*. Among the 26 predicted TRs, Ctnnb1^39^ and Srebf1^40^ have been reported as regulators of *Scd2*, while *Nr1h3* and *Scd2* are co-regulated in the PPAR signaling pathway^41^, which plays a critical role in fetal liver by regulating lipid metabolism^42^. In addition, the KnockTF database^43^ reported *Scd2* as differentially expressed after either Nrf1 or Smad4 loss-of-function treatment (knock-down or knock-out), indicating that Nrf1 and Smad4’s regulatory role to *Scd2*. Therefore, Dory can help identify promoter-enhancer interactions *de novo* from specifically designed high-resolution chromatin tracing data.

### Dory identifies potential regulators during mouse lung cancer progression

Finally, we sought to apply Dory to study chromatin alterations during disease progression. We analyzed the chromatin tracing data from a lung cancer mouse model that includes preinvasive cancer cells (lung adenoma) and invasive cancer cells (lung adenocarcinoma, LUAD)^20,32^. In this dataset, genomic regions spanning 19 preidentified candidate progression driver (CPD) genes in adenoma and LUAD were probed using chromatin tracing. For each CPD, 40 genomic regions spanning the CPD promoter and adjacent potential cis-regulatory regions were traced at a resolution ranging from 5kb to 20kb. We calculated the pairwise distances for each CPD’s trace and found that the region pairs with the same genomic distance exhibit shorter spatial distances in LUAD than in adenoma for every CPD, indicating a significant trend of chromatin compaction from adenoma to LUAD at this fine scale (Fig. 5a, Supplementary Fig. 6), consistent with prior analyses^20^. To correct the overall compaction effect, we normalized the pairwise distance matrix by the average value of each matrix for all chromatin traces for the CPD (See details in Methods). Then we applied Dory to the normalized distance matrices in LUAD and adenoma to generate the DiffScore matrix for every CPD. We identified CloserRegions whose distance to the CPD promoter is significantly shorter in LUAD (invasive cancer cells) than in adenoma (preinvasive cancer cells) (Fig. 5b, c, Supplementary Fig. 7, 8). To further examine whether the identified CloserRegions (LUAD vs. adenoma) are likely to interact with the CPD promoter in LUAD, we surveyed the spatial distance of all region pairs at the same genomic distance as the CloserRegion to the CPD promoter in LUAD, and found that the spatial distance between the CloserRegion and the promoter is significantly shorter than others (Supplementary Fig. 9a). These indicate CloserRegion is more likely to contain an enhancer of the CPD than other probed regions. We analyzed all 19 CPD traces using the same approach and identified the CloserRegions in the traces for CPDs *Mycn*, *Myc*, *Anxa8*, *Fhit*, and *Oit1* (Supplementary Fig. 7, 8, 9b-e).

**Figure 5.**
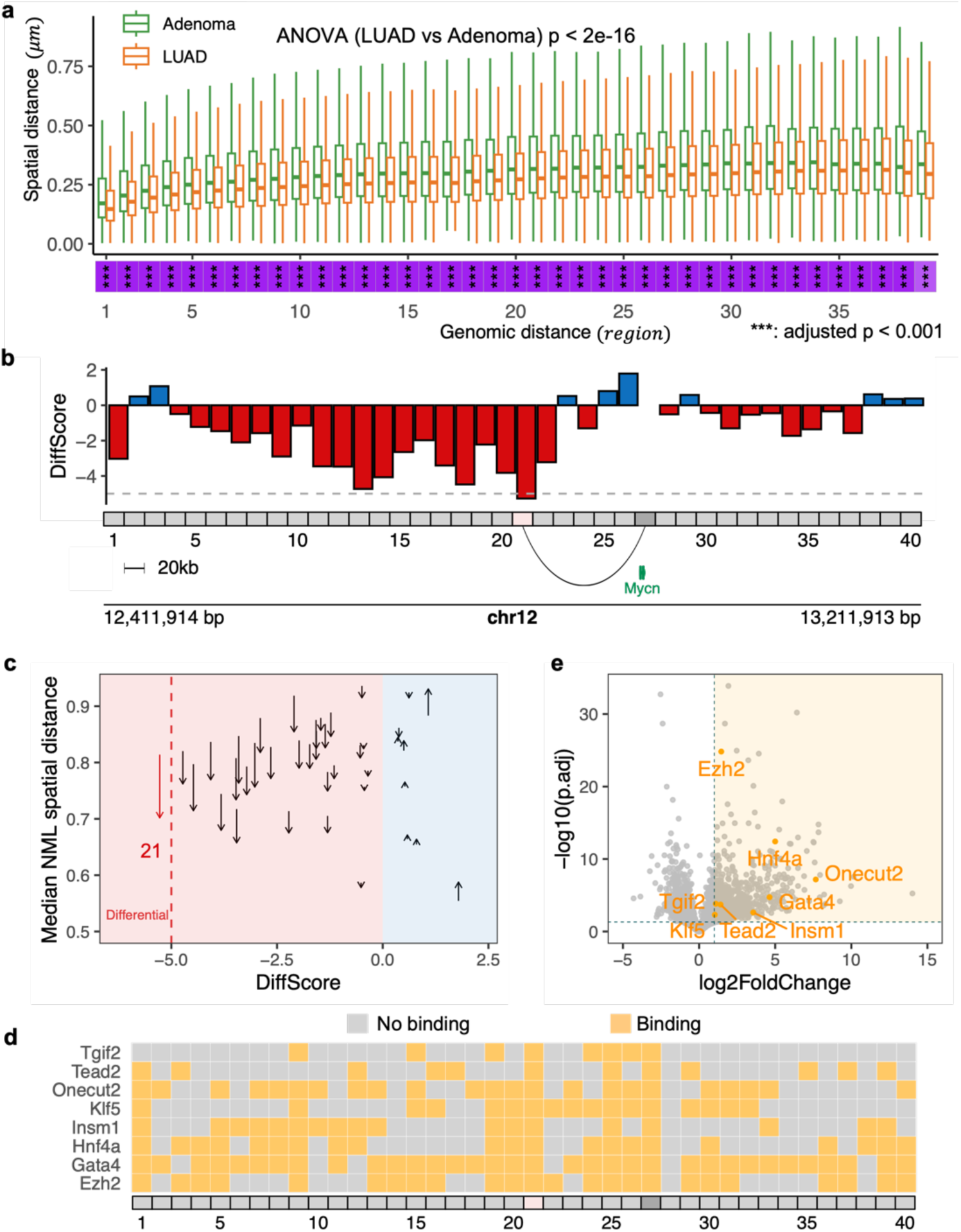
Dory identifies transcriptional regulators during mouse lung cancer progression. **a**, Box plots displaying the spatial distance (y-axis) between region pairs across different genomic distance (x-axis) in adenoma (green) and LUAD (orange). Data are from 40 probed genomic regions spanning the *Mycn* promoter. **b**, Bar plot displaying the DiffScores for region pairs of *Mycn* promoter (Region 27, indicated by dark gray box) and 39 flanking genomic regions comparing LUAD with adenoma. Region 21 is identified as a CloserRegion (pink box). **c**, Arrow plot displaying normalized (NML) spatial distance patterns between the *Mycn* promoter and the 39 flanking regions from adenoma to LUAD. The x-axis represents the DiffScore. The y-axis represents the median NML spatial distance across all traces. Each region pair is represented as an arrow from adenoma (arrow start) to LUAD (arrow end). The red dashed line indicates the DiffScore threshold. The red arrow indicates the CloserRegion. **d**, Regulator (y-axis) binding sites on CloserRegions (pink region on x-axis). Orange boxes indicate binding; gray boxes indicate no binding. **e**, Volcano plot showing the differential gene expression between LUAD and adenoma. The shaded region highlights genes significantly upregulated in LUAD than in adenoma (log2FoldChange > 1 & p.adj < 0.05). Orange dots indicate the TR genes that potentially bind to the CloserRegion (Region 21) in the *Mycn* trace.

To identify the potential regulators of the CPDs in LUAD, we then utilized the ChIP-seq profiles collected in the BART library^36^ to identify the TRs bound in the CloserRegions for each CPD. We further narrowed down the list of potential regulators by requiring the differential expression^20^ of the regulators in LUAD compared to adenoma. As a result, we identified Ezh2, Gata4, Hnf4a, Insm1, Klf5, Onecut2, Tead2 and Tgif2 as regulators of *Mycn* (Fig. 5d, e), as well as potential regulators for other CPDs *Myc*, *Anxa8*, *Fhit*, and *Oit1* (Supplementary Fig. 10-11).

Taken together, Dory helps to investigate chromatin alterations during cancer progression and provides valuable perspectives on identifying active enhancers and candidate transcriptional regulators of progression driver genes.

## Discussion

Dory is an efficient computational tool for differential analysis of image-based chromatin tracing data. We assessed Dory’s performance and observed the high robustness and high reproducibility in quantifying differential patterns between chromatin tracing samples using a subsampling approach. We applied Dory to a chromatin tracing dataset for chromosome 19 across six cell types from mouse fetal liver and showed that large-scale chromatin structure changes identified by Dory are partially associated with A/B compartment changes. From two case studies for high-resolution (5kb) chromatin tracing in a mouse fetal liver sample and a lung cancer progression model, respectively, we demonstrated that Dory can identify active enhancer regions and potential transcriptional regulators of genes of interest.

One unique feature of Dory is the ability to independently test each pair of genomic regions from a chromatin tracing dataset. This direct approach enables quantitative comparison without requiring chromatin structure reconstruction, which is often computationally complex and prone to over-fitting. By leveraging the statistical properties of unpaired tests, Dory also circumvents issues associated with missing data, a common challenge in chromatin tracing experiments. This straightforward design makes Dory both versatile and computationally efficient. Furthermore, Dory supports multi-group chromatin tracing datasets that allows for pairwise comparisons among multiple cell types or experimental groups, beyond the typical two-state analyses used in most existing methods.

Dory is a distinctive tool that provides functionalities unavailable in other existing methods. For example, while SnapFISH^30^ is designed to identify chromatin loops from chromatin tracing data within a single biological condition, it lacks the ability to perform differential analysis between two or more groups of chromatin traces. In contrast, Dory is specifically developed to detect region pairs exhibiting significantly different spatial distances across biological conditions, regardless of whether these region pairs form chromatin loops. To identify differential chromatin loops, users can combine the strength of SnapFISH for loop calling and Dory for differential analysis when analyzing high-resolution chromatin tracing data. Furthermore, while SnapFISH employs a two-sample t-test, Dory adopts the Wilcoxon rank-sum test as its default statistical framework for comparing distance distributions two conditions. This choice is motivated by the observation that spatial distances of region pairs across chromatin traces deviate from a normal distribution, as confirmed by our normality check using the K-S test (Supplementary Fig. 12). Therefore, a non-parametric test is implemented as the default to ensure robustness in statistical inference.

While analyzing chromatin tracing data with Dory provides valuable insights into 3D genome organization, it is also important to recognize the limitations of Dory and potential directions for future improvements. As a fully data-driven framework, Dory’s accuracy is constrained by the resolution of the image-based chromatin tracing data, which is determined at the experimental design stage. Moreover, Dory assumes independence among genomic regions; potential correlations between neighboring regions or regions within the same TADs are not explicitly considered in the model, even though such associations can be biologically meaningful depending on resolution and context. Currently, Dory does not include intrachromosomal interactions, primarily due to computing complexity considerations, although this is an interesting direction for future explorations^44,45^. Additionally, while Dory’s statistical framework can automatically handle datasets with missing values, the presence of missing data inevitably reduces statistical power. Various strategies for missing data imputation have been proposed, such as filling missing region-pair distances with the mean or median across all chromatin traces^30^, or applying model-based approaches such as SnapFISH-IMPUTE^31^. However, such imputation often reduces data variability, increases the risk of type I error (false positives), and significantly decreases computational efficiency. Developing more effective computational approaches to handle missing data remains a key challenge for future research.

## Methods

### Spatial distance calculation

The input of Dory is two sets of chromatin traces, following the 4DN format^32^. Each chromatin trace consists of the 3D coordinates of the probed genomic regions. For the 3D coordinates (*x*_*i*_, *y*_*i*_, *z*_*i*_) for genomic region *i* and (*x*_*j*_, *y*_*j*_, *z*_*j*_) for genomic region *j*, we calculated the spatial distance between the region pair (*i*, *j*) using the Euclidean distance formula as 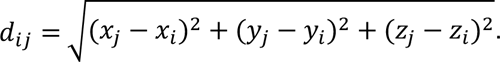

### Statistical test in Dory algorithm

For *M* chromatin traces in Condition 1 and *N* chromatin traces in Condition 2, the distance matrices are represented by *D*_1,*m*_ and *D*_2,*n*_, in which *m* ∈ (1,2, …, *M*) and *n* ∈ (1,2, …, *N*). We utilized the Wilcoxon rank sum test to do the differential analysis of two spatial distance vectors of a region pair corresponding to two conditions. In Condition 1, the spatial distance between region pair (*i*, *j*) across *M* chromatin traces is denoted by the vector *D*_1*i*j_ = (*d*_1*ij*,1_, *d*_1*ij*,2_, …, *d*_1*ij*,*m*_, …, *d*_1*ij*,*M*_); in Condition 2, the spatial distance between region pair (*i*, *i*) across *N* chromatin traces is denoted by the vector *D*_2*i*j_ = (*d*_2*ij*,1_, *d*_2*ij*,2_, …, *d*_2*ij*,*n*_, …, *d*_2*ij*,*N*_). For *d*_2*i*j,1_, 2: Condition 2; *i*: region *i*; *j*: region *j*; 1: the 1^st^ chromatin trace in this condition. We calculated the difference of spatial distance of this region pair using

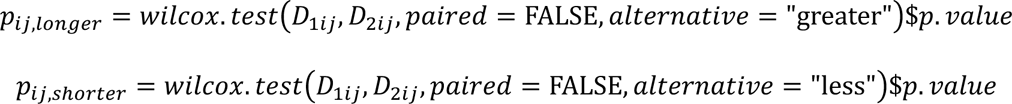

in R. Then, the DiffScore for this region pair was defined as

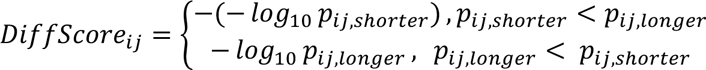

We also examined the two-sample t-test on the spatial distance vectors. In the two-sample t- test, it assumes that the data follows the normal distribution. To investigate whether the spatial distance data follows a normal distribution, we utilized the K-S test to assess its normality. The adjusted p values for the tests are around 1e-6 (Supplementary Fig. 12), which rejects the null hypothesis that the data follows the normal distribution. However, we noticed that after logarithmically transform the spatial distance vector, the adjusted p values for the normalization test increase to around 0.59 (the -log10(adj. pvalue) decreases), which is larger than 0.05.(Supplementary Fig. 12). We could not reject the hypothesis that the logarithmically transform distance data follow the normal distribution, that is, the data distribution tends to be normal. Therefore, we recommended the two-sample t-test for log-transformed not the original spatial distance data for chromatin tracing.

### Missing data processing

In the *wilcox*. *test*, the parameter *paired* was set to be FALSE which allowed the unpaired comparison, that is, the vector length of *D*_1*i*j_ and *D*_2*i*j_ could be different. If there were missing coordinates for regions leading to the missing spatial distance for region pairs, we just left these distances as *NAs*, which still could be processed by the *wilcox*. *test* function.

### Parallel analysis for multiple region pairs

For each region pair, we implemented a Wilcoxon rank sum test. To accelerate the DiffScore calculation across thousands of region pairs with thousands of test iterations, we utilized the *future* and *furrr* packages in R to perform the parallel computation.

### Output of Dory

For a series of genomic regions in the comparison of condition 2 VS condition 1, the output of Dory includes 1) the spatial distance of region pairs in each condition, 2) the DiffScore matrix for region pairs and 3) Differential region pairs from greater and less direction.

### Chromatin tracing dataset in mouse fetal liver

We collected chromatin tracing data in mouse fetal liver from 4DN data portal^15,32^. Two datasets were included. One included the chromatin traces of 50 genomic regions along the chromosome 19 in six cell types: proerythroblasts, erythroblasts, hepatocytes, macrophages, megakaryocytes, and endothelial cells. The other included the chromatin traces of 19 genomic regions upstream of Scd2 promoter with genomic resolution of 5kb in these six cell types.

### Subsampling of chromatin traces

For the 1490 chromatin traces in proerythroblasts and 2615 chromatin traces in erythroblasts, we separately subsampled 100, 200, …, 1000 traces from each cell type. To reduce the selection bias in random sampling, we performed 20 independent random samplings for each subsampling scenario. For example, when selecting 100 traces, we generated 20 random cases, each involving the selection of 100 chromatin traces from the entire chromatin traces.

### Calculation of Pearson correlation of DiffScore between subsamples and the whole sample

We applied Dory to the whole chromatin traces in erythroblast and proerythroblast and generated DiffScores for spatial distance of region pairs. For each subsampling scenario, we applied Dory to each randomly selected case to generate the DiffScores. Finally, we compared the PCC of DiffScores generated from using the randomly selected subsamples and from using the whole traces. We also utilized the *cor*. *test* in R to assess the statistical significance of the PCC.

### Calculation of reproducibility rate

For each subsampling scenario, we performed 20 random samplings. To measure the reproducibility of Dory across these random samples with the same number of chromatin traces, we defined the reproducibility rate in two directions: distance increasing (becoming longer) and distance decreasing (becoming shorter). For each random sampling, the reproducibility rate in a given direction is defined as the percentage of top 50 differential region pairs from that sample also appeared in the top 50 differential region pairs of any of the other 19 random samplings.

### Definition of ABchange score

For each region, we examined its A/B compartment score (*C*) with the same computational workflow for Hi-C data in previous studies^10,15^. We then calculated the A/B compartment interactions of a region pair (*i*, *j*) by multiplying the compartment scores of this region pair as *C*_*i*_ ∗ *C*_*j*_. Lastly, from Condition 1 to Condition 2, the ABchange is defined as the difference in the interaction score of this region pair, calculated as *C*_2,*i*_ ∗ *C*_2,j_ − *C*_1,*i*_ ∗ *C*_1,j_ (Fig. 3a).

### Calculation of PCC between DiffScore and ABchange

For the comparison of two conditions, we calculated the PCC between DiffScore generated by Dory and ABchange scores. We utilized the *cor*. *test* in R to assess the statistical significance of the PCC.

We also implemented the permutation for the ABchange scores. We randomly shuffled the ABchange scores 1000 times and recalculated the PCCs between DiffScore and ABchange for each permutation. This PCC set served as a control for the PCC between real ABchange and DiffScore.

### Permutation test for the correlation between DiffScore and ABchange

For the comparison of two cell types, we permuted the ABchange scores1000 times to generate 1000 ‘background’ ABchange vectors. Then we recalculated the PCC between DiffScore and every ‘background’ ABchange vector to generate the ‘background’ PCC for every permutated background.

### Identification of outliers in the correlation between DiffScore and ABchange

In the linear relationship between DiffScore and ABchange, we identified the outliers by the residual and correlation coefficient. First, we identified the dots with large residuals by |*residual*| > 3 ∗ *SD*_*residual*_ (*SD*: standard deviation). Then, we examined the correlation coefficient after removing each large-residual dot. If the absolute value of the correlation coefficient increases after removing this dot, then the dot was identified as an outlier (orange dots in Fig. 3b and Supplementary Fig. 3. Each orange dot represents an outlier region pair).

### Gene expression profiles in mouse fetal liver

We downloaded the single-cell digital gene expression matrix for fetal liver from the Mouse Cell Atlas^38^. We obtained the read count for 12568 genes in each cell with cell type labels. We selected cells labeled as hepatocyte, macrophage, megakaryocyte, proerythroblast, and erythroblast (no endothelial cells are identified in this single-cell expression dataset). Then we utilized the NormalizeData in Seurat^46^ to get the normalized expression of genes in each cell type. The normalization was performed as 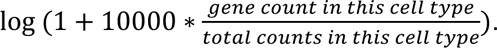 We utilized FindMarkers in Seurat to identify the differentially expressed genes between any two cell types.

### Fisher’s exact test for up-regulated genes on shorter outlier regions

For the comparison of each pair of cell types, we collected the outlier region pairs in the relationship between DiffScore and ABchange. It is important to note that for any given pair of cell types, like A and B, the outliers in the shorter direction for the A-to-B comparison are the outliers in the longer direction for the B-to-A comparison. To avoid the redundancy, for all cell-type-pairs’ comparisons, including A-to-B and B-to-A, we only consider the outliers in the shorter direction.

Next, for each pair of cell types, we selected the unique shorter outlier regions from the set of outlier region pairs. We emphasize ‘unique’ because, since outliers were defined as region pairs, some regions could simultaneously appear in both longer and shorter outliers. To ensure clarity, we excluded these overlapping regions from the set of shorter outlier regions.

Finally, we counted the up-regulated genes and down-regulated genes located in shorter direction and non-shorter direction and aggregated these counts across all pairs of cell types. We then performed the Fisher’s exact test using

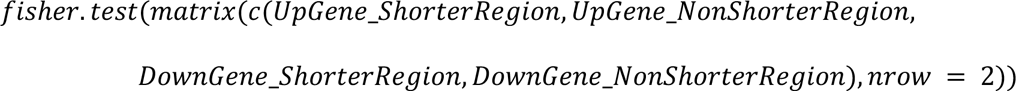

in R to get the odds ratio and p-value.

### Identification of regulators’ binding sites on DiffRegions

In BART library^36^, it utilizes the union DNaseI hypersensitive sites (UDHS) as a platform to save the regulators’ binding sites based on ChIP-seq profiles in mouse. For each UDHS, the BART library contains the regulators binding on that region. We first identified the UDHS that overlapped with DiffRegions^47^. Then, we collected the regulators those bind on the identified UDHS set.

### Chromatin tracing dataset in lung cancer progression

We collected chromatin tracing data for genomic regions surrounding candidate progression driver (CPD) genes in a mouse model of lung cancer progression^20^, from adenoma to lung adenocarcinoma (LUAD). For each CPD, 40 consecutive genomic regions spanning its promoter were traced in both adenoma and LUAD cells, with genomic resolutions ranging from 5kb to 20kb.

### Normalization of spatial distance in adenoma and LUAD

To normalize the spatial distance of region pairs, we calculated a normalization factor for each state. For every distance matrix in a given state, we first calculated the mean of all spatial distances in this distance matrix, denoted as *E*(*Trace*). We then calculated the mean across all traces in this state, denoted as *E*(*E*(*Trace*)), which serves as the normalization factor. We normalized each element in the distance matrix across all traces by dividing it by this normalization factor to remove the overall compaction/decompaction effects.

### Differential gene expression profiles in adenoma and LUAD

We downloaded differential gene expression profiles from the lung cancer progression study^20^. It includes the log2 fold change and adjusted p-value for each gene in the comparison of LUAD to adenoma.

## Code availability

Dory is implemented in R and the open-source package is available at: https://github.com/zang-lab/Dory

## Competing interests

Si.W. and M.L. are inventors on a patent applied for by Yale University related to cancer 3D genome mapping by chromatin tracing. Si.W. is an inventor on patent applications related to the chromatin tracing technology.

## Acknowledgements

We thank members of the Wang Lab (especially Mengwei Hu) and Zang Lab for helpful discussions. This work was partially supported by NIH grants R35GM133712 (C.Z.), R01CA292936 (Si.W.), UH3CA268202 (Si.W.), and R01HG013503 (Si.W.), as well as Cancer Center Support Grant (C.Z.) and Farrow Fellowship (Z.M.) within the University of Virginia Comprehensive Cancer Center.

## Notes

### Competing Interest Statement

Siyuan Wang and Miao Liu are inventors on a patent applied for by Yale University related to cancer 3D genome mapping by chromatin tracing. Siyuan Wang is an inventor on patent applications related to the chromatin tracing technology.

