## Supplementary Figures for "Differential analysis of image-based chromatin tracing data with Dory"

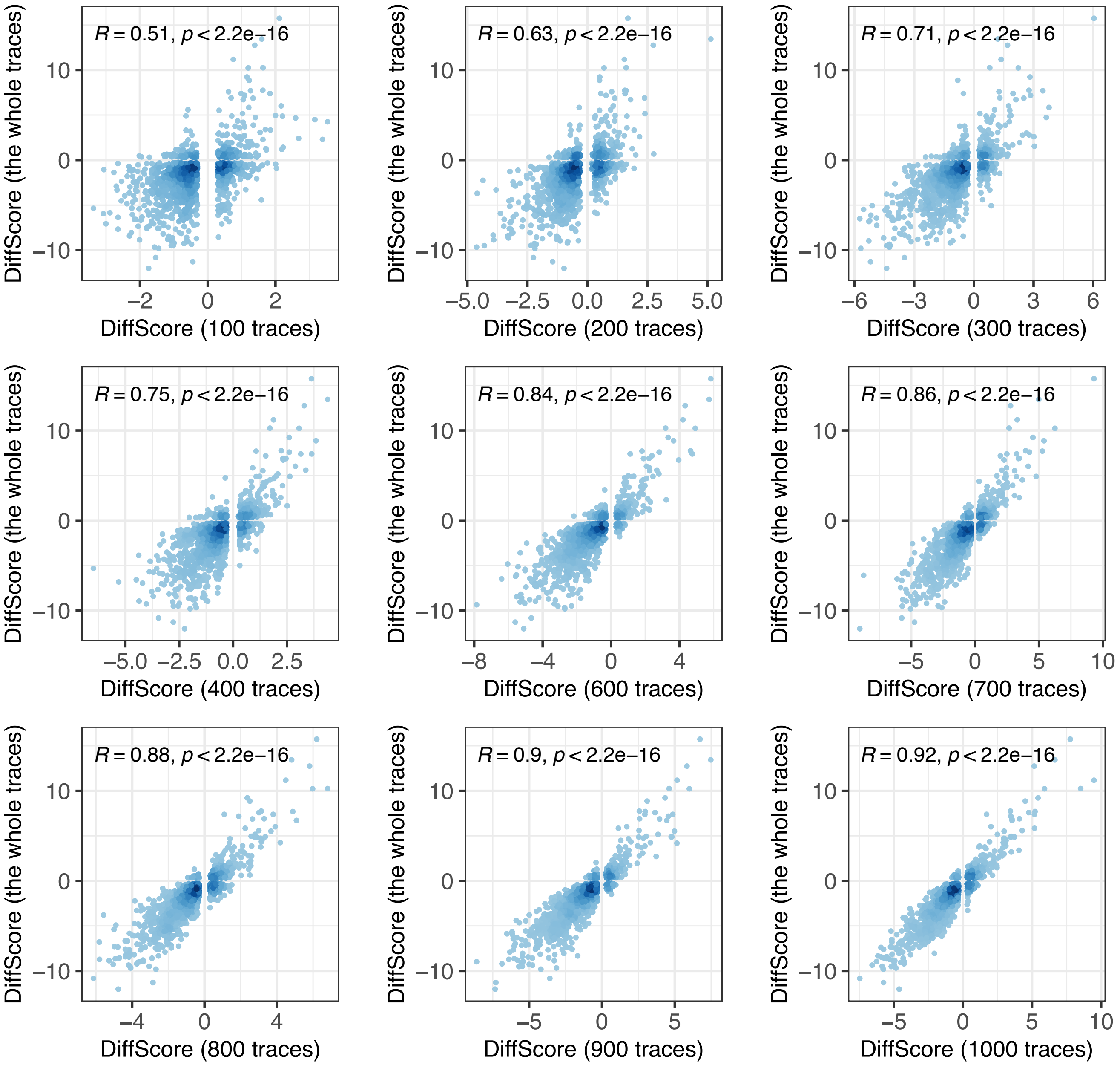


**Supplementary Fig. 1. Scatter plots displaying the correlation of DiffScore calculated from the whole samples and the subsamples (100, 200, ... 1000 traces for each cell type) for the comparison of proerythroblast and erythroblast.** For every sub-trace sample, only one case from 20 random samples is selected.

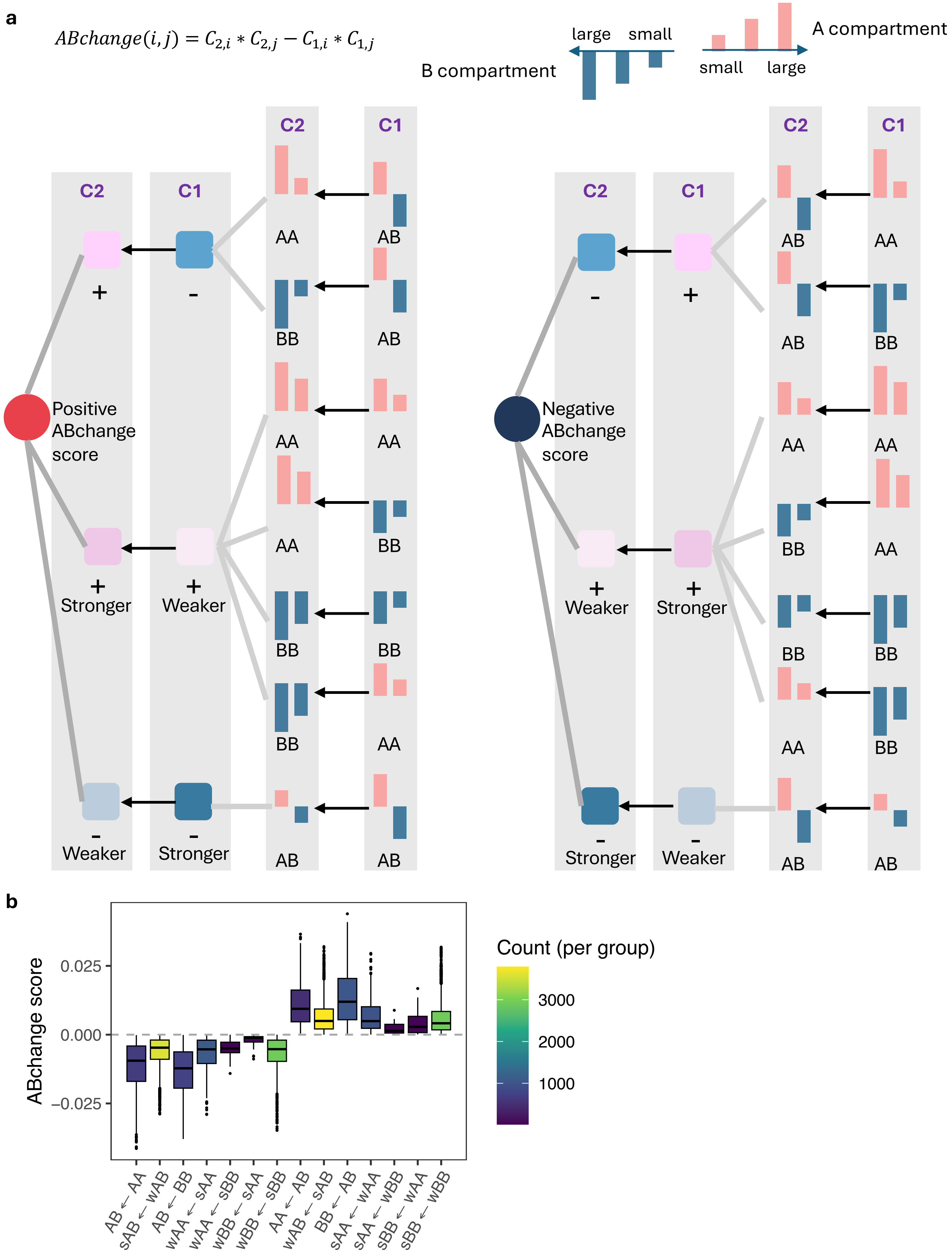

a
$ABchange(i,j) = C_{2,i} * C_{2,j} - C_{1,i} * C_{1,j}$
large
small
B compartment
small
large
A compartment
C2
C1
AA
AB
BB
Positive ABchange score
Negative ABchange score
+
-
Stronger
Weaker
b
ABchange score
0.025
0.000
−0.025
AB ← AA
sAB ← wAB
AB ← BB
wAA ← sAA
wAA ← sBB
wBB ← sAA
wBB ← sBB
AA ← AB
wAB ← sAB
BB ← AB
sAA ← wAA
sAA ← wBB
sBB ← wAA
sBB ← wBB
Count (per group)
3000
2000
1000

**Supplementary Fig. 2. Schematic interpretation of A/B compartment score, A/B interaction score, and ABchange score.** A/B change score is calculated by A/B interaction score in Condition 2 minus in Condition 1. A/B interaction score in every condition is the product of compartment scores for the two regions. **a**, Left panel: positive ABchange derived from three types of A/B interaction score (positive – negative, stronger positive – weaker positive, and weaker negative – stronger negative), which is originally from seven types of A/B compartment score (AA - AB, BB – AB, stronger AA – weaker AA, stronger AA – weaker BB, stronger BB – weaker BB, stronger BB – weaker AA, and weaker AB – stronger AB). Right panel: negative ABchange derived from three types of A/B interaction score (negative – positive, weaker positive – stronger positive, and stronger negative – weaker negative), which is originally from seven types of A/B compartment score (AB - AA, AB – BB, weaker AA – stronger AA, weaker BB – stronger AA, weaker BB – stronger AA, weaker AA – stronger BB, and stronger AB – weaker AB). **b,** box plot displaying the distribution of ABchange in every type of A/B compartment change. The box color represents the number of data points in each group/box.

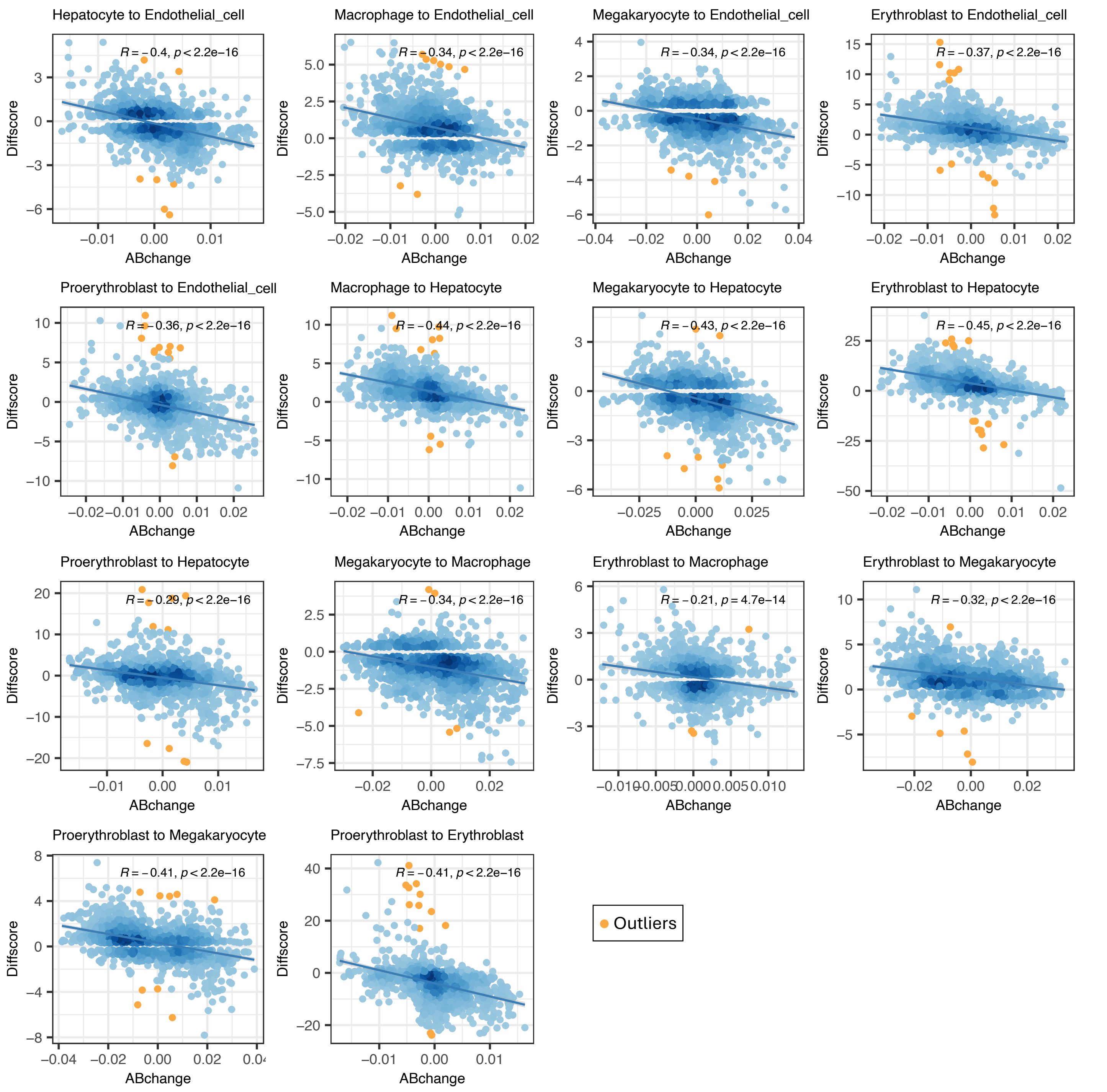


**Supplementary Fig. 3. Scatter plots displaying the correlation between the DiffScore and ABchange in the comparison of two cell types** (the panel title: Condition 1 to Condition 2. i.e., in the last panel, Proerythroblast is Condition1; Erythroblast is Condition 2). The outliers are highlighted in orange. The darkness of blue dots indicates the dots density.

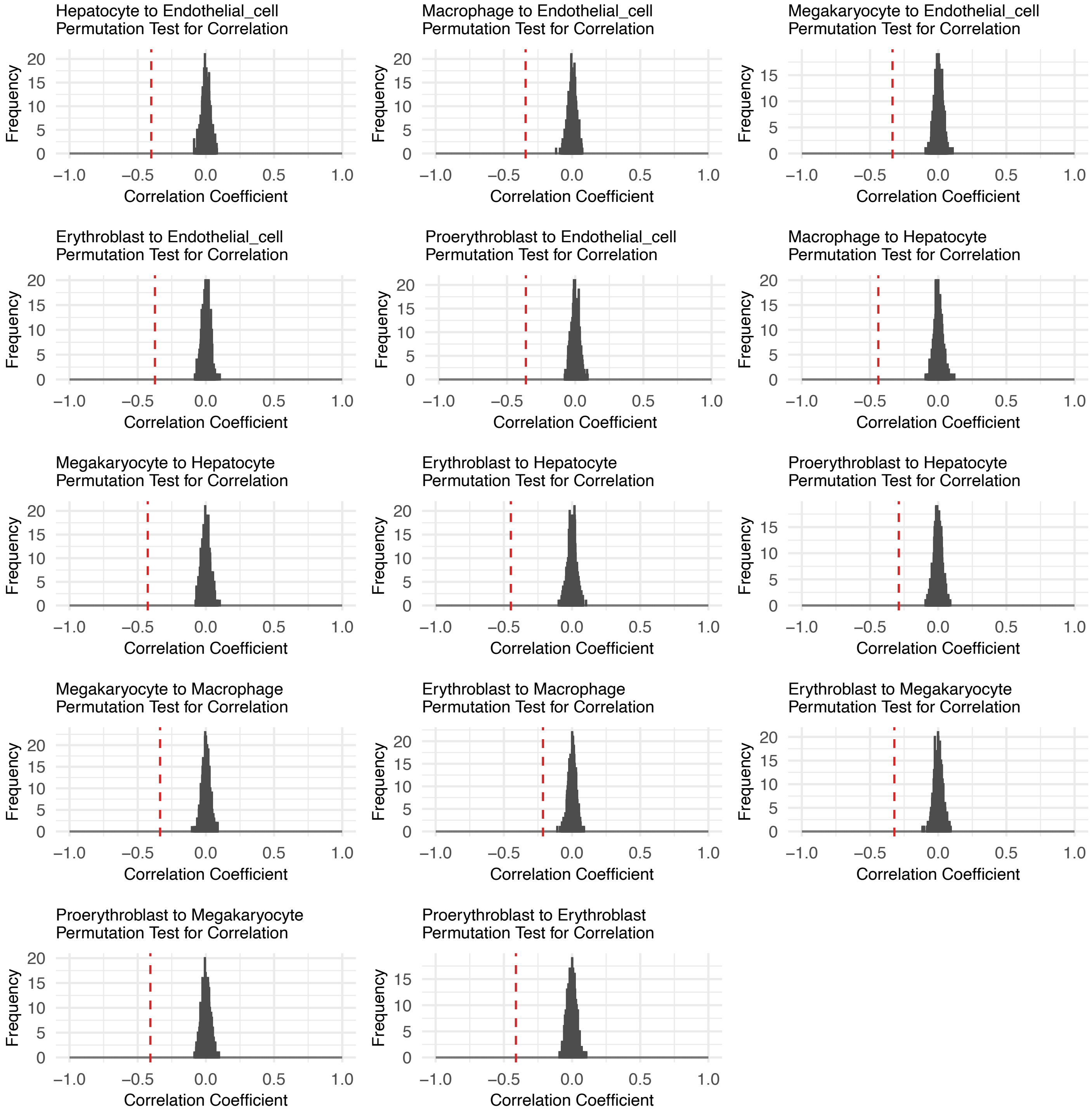


**Supplementary Fig. 4.** Permutation test for the correlation between DiffScore and ABchange in the comparison of other cell types. The null distribution (around 0) is the correlations from 1000 samples of randomly permuted ABchanges. The red dashed line indicates the observed correlation coefficient between DiffScore and ABchange (in Supplementary Figure 3).

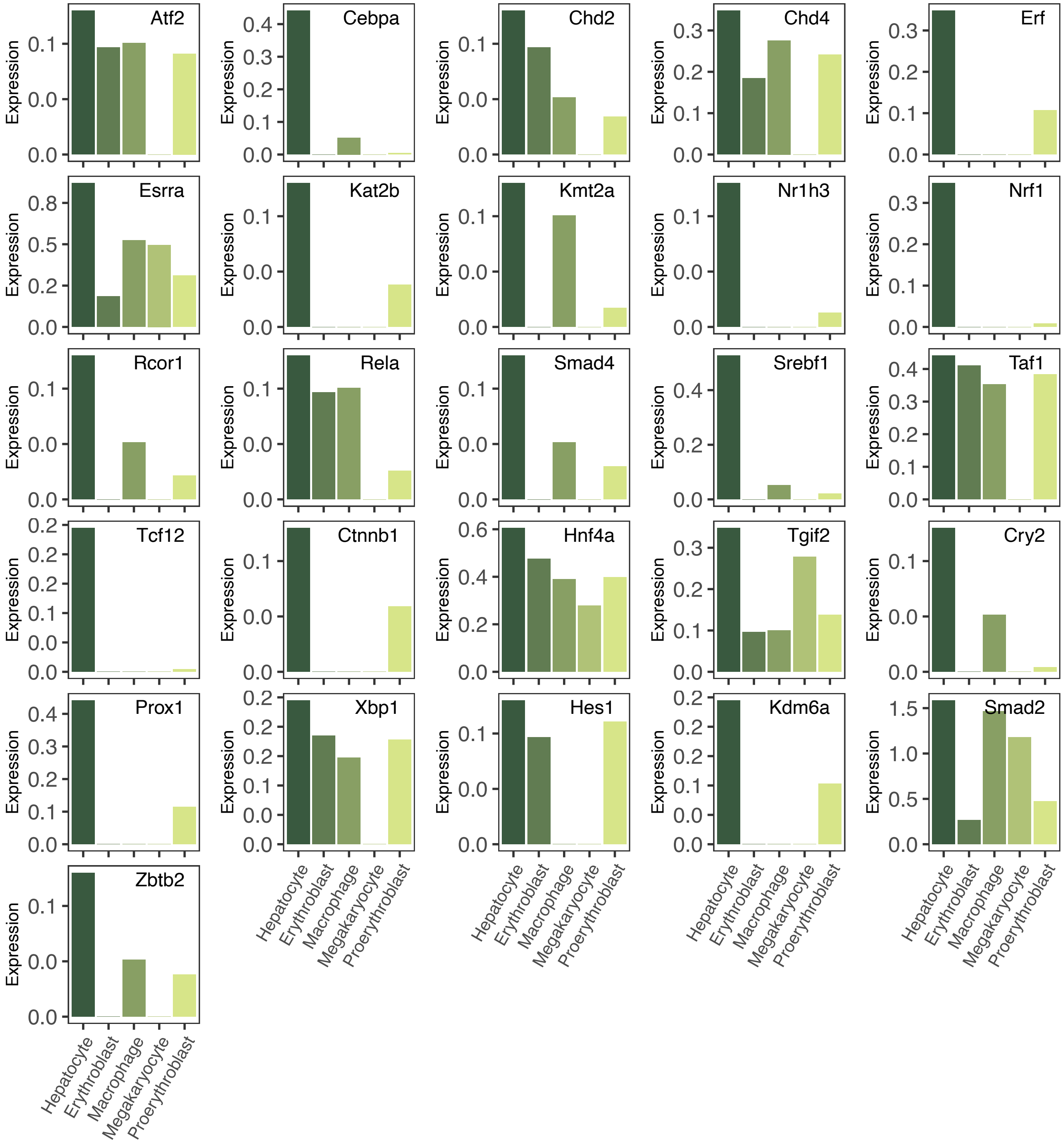


**Supplementary Fig. 5. Normalized expression levels of identified potential regulators of Scd2 in hepatocytes and four other cell types in mouse fetal liver from the Mouse Cell Atlas.**

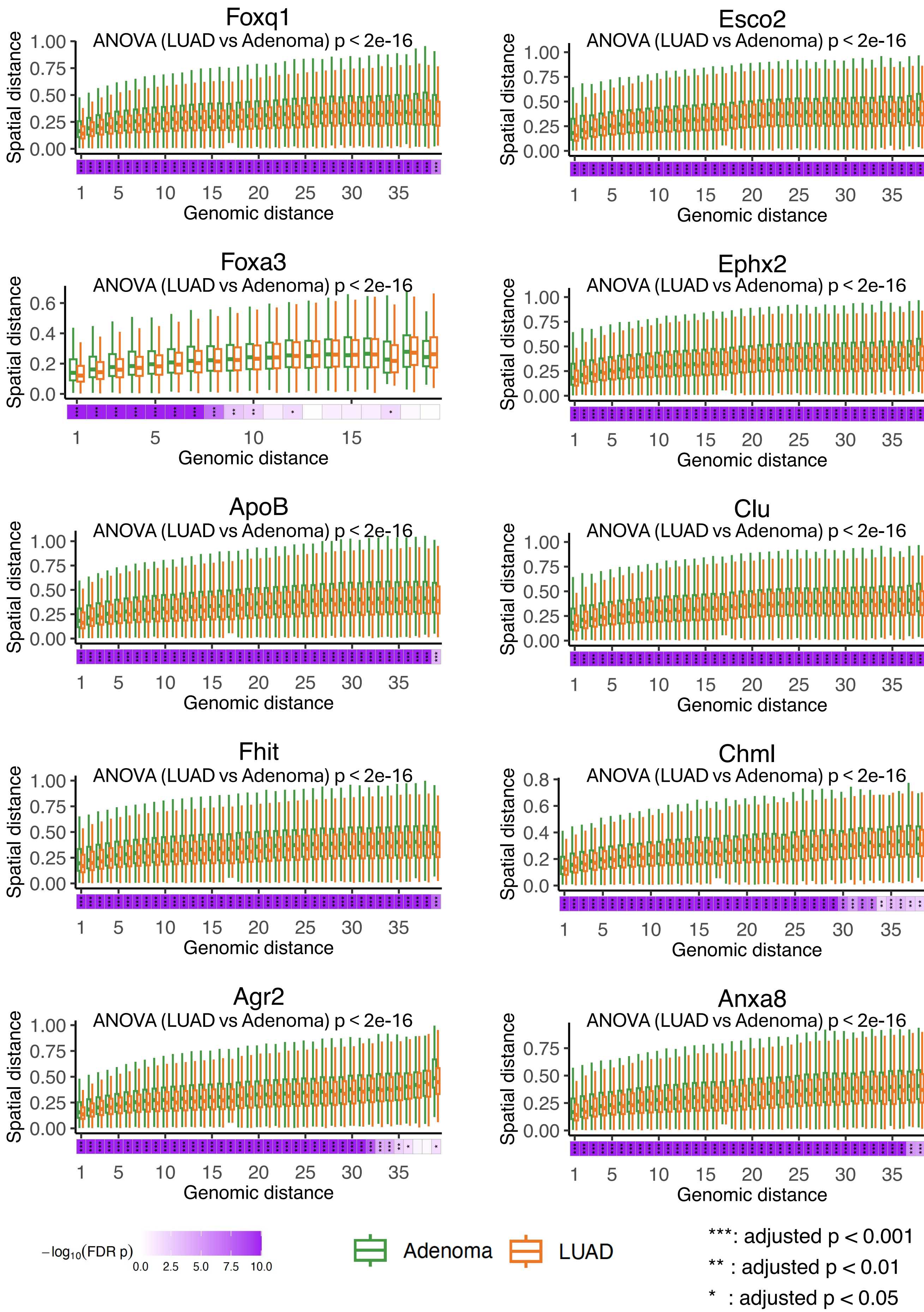

Foxq1
ANOVA (LUAD vs Adenoma) p < 2e-16
Esco2
ANOVA (LUAD vs Adenoma) p < 2e-16
Foxa3
ANOVA (LUAD vs Adenoma) p < 2e-16
Ephx2
ANOVA (LUAD vs Adenoma) p < 2e-16
ApoB
ANOVA (LUAD vs Adenoma) p < 2e-16
Clu
ANOVA (LUAD vs Adenoma) p < 2e-16
Fhit
ANOVA (LUAD vs Adenoma) p < 2e-16
Chml
ANOVA (LUAD vs Adenoma) p < 2e-16
Agr2
ANOVA (LUAD vs Adenoma) p < 2e-16
Anxa8
ANOVA (LUAD vs Adenoma) p < 2e-16
Spatial distance
Genomic distance
$-\log_{10}$(FDR p)
0.0 2.5 5.0 7.5 10.0
Adenoma
LUAD
***: adjusted p < 0.001
** : adjusted p < 0.01
* : adjusted p < 0.05

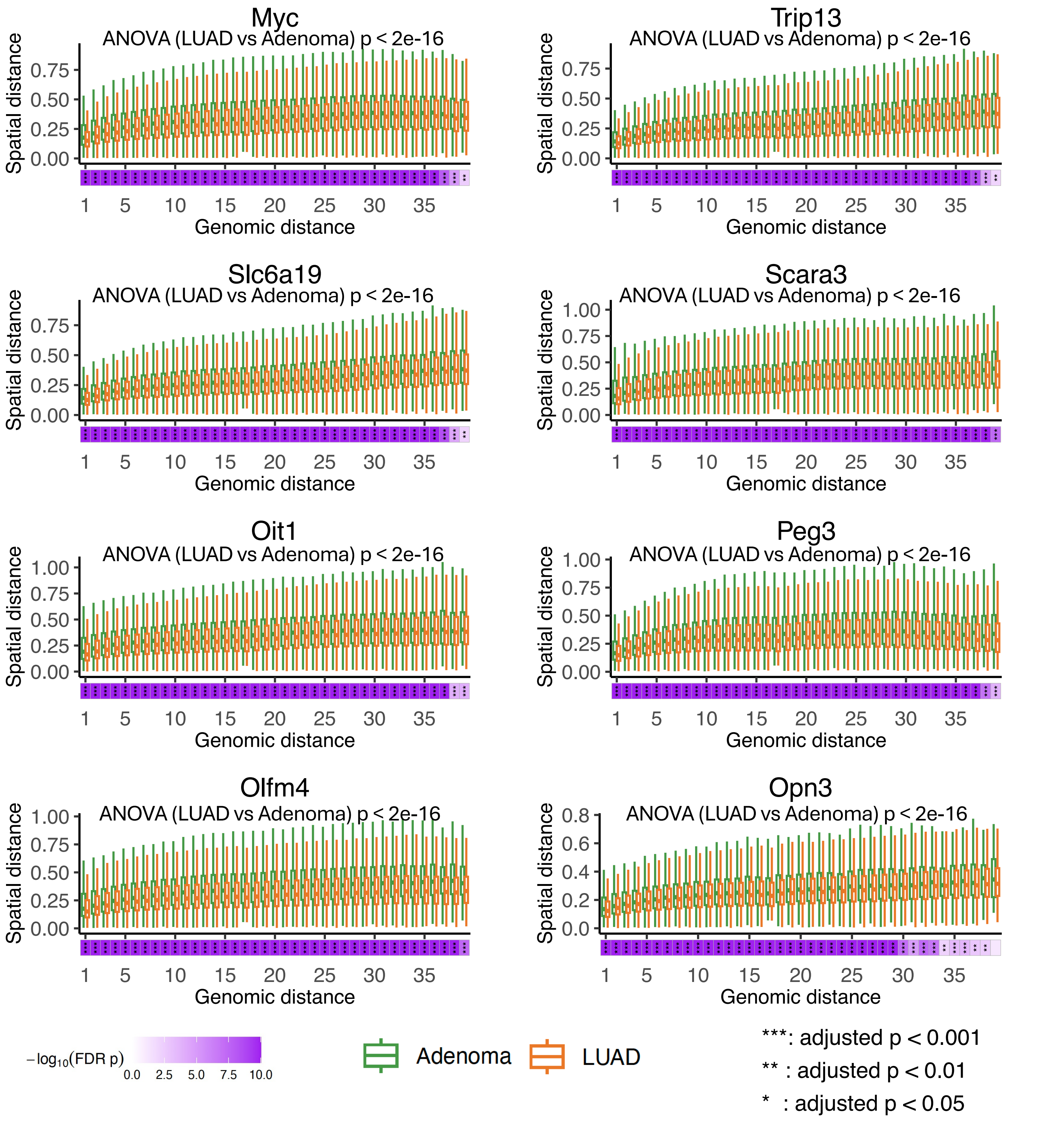


**Supplementary Fig. 6. Box plots displaying the spatial distance (y-axis) between region pairs across different genomic distance (x-axis) in adenoma (green) and LUAD (orange) for 18 other CPDs' trace.** The ANOVA test for LUAD verse adenoma displaying a significant difference of spatial distance. The purple bar and star annotation (*) indicate the significance of difference for region pairs in every genomic distance.

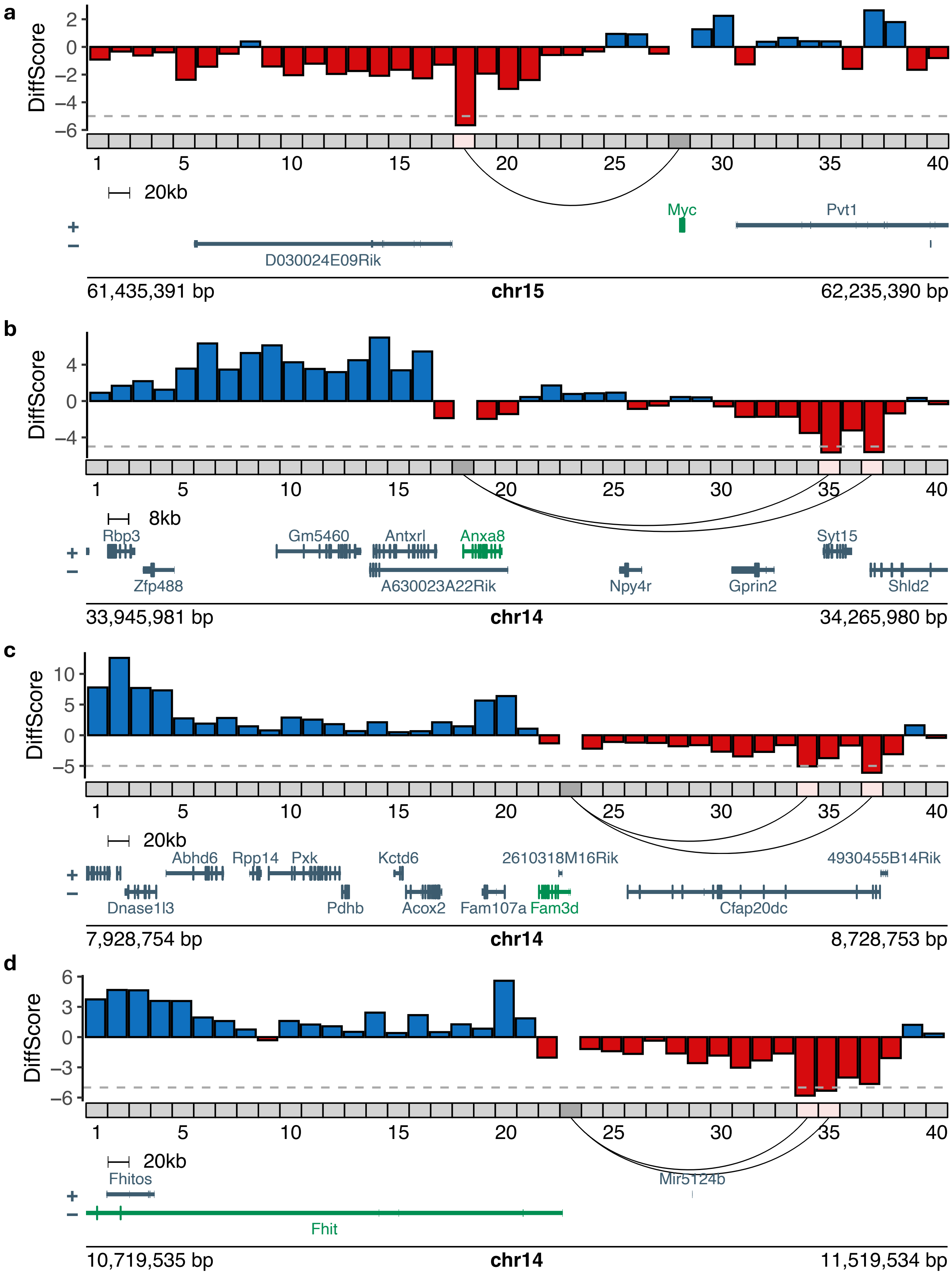


**Supplementary Fig. 7. Bar plot displaying the DiffScores for region pairs of CPD promoter (dark gray) and 39 genomic regions (light gray) spanning the promoter comparing LUAD with adenoma. a,** for *Myc*. **b,** for *Anxa8*. **c,** for *Oit1*. (note: Oit1 is also known as Fam3d) **d,** for *Fhit.* The pink regions are identified as CloserRegions.

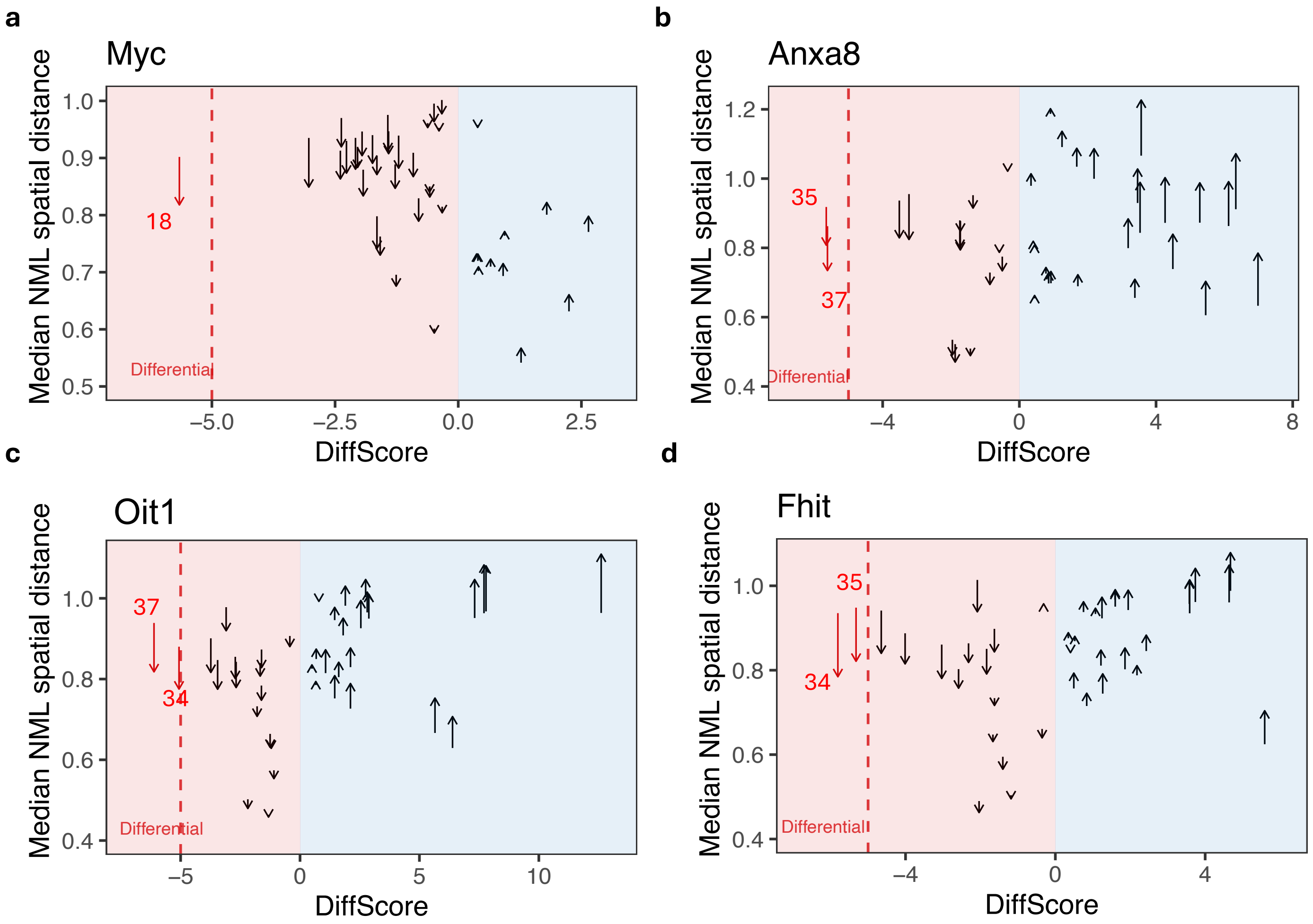


**Supplementary Fig. 8. Arrow plot displaying normalized (NML) spatial distance patterns between the CPD promoter and 39 flanking regions.** Each region pair is represented as an arrow from adenoma (arrow start) to LUAD (arrow end). The CloserRegions are labeled by red text and red arrows. **a,** for *Myc*. **b,** for *Anxa8*. **c,** for *Oit1*. **d,** for *Fhit.*

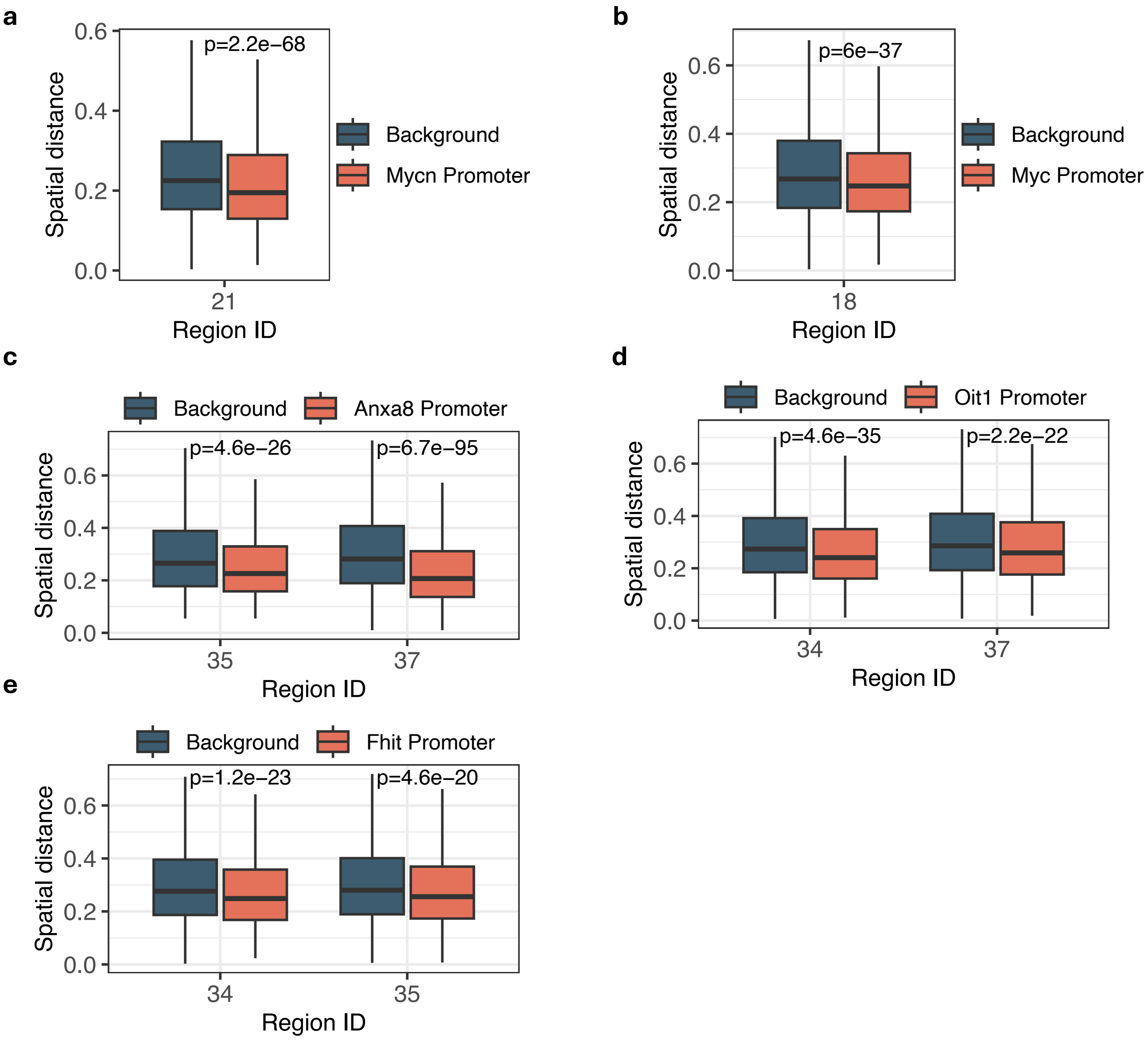


**Supplementary Fig. 9. Box plots displaying the spatial distance (y-axis) between region pairs with the same genomic distance as CloserRegion (ID in x-axis) and the CPD promoter.** Red box represents the spatial distance between the CloserRegion and the CPD promoter. Blue box represents the spatial distance between region pairs with the same genomic distance as the CloserRegion and the CPD promoter. **a,** CloserRegion (Region 21) for *Mycn*. **b,** CloserRegion (Region 18) for *Myc*. **c,** CloserRegions (Regions 35 and 37) for *Anxa8*. **d,** CloserRegions (Regions 34 and 37) for *Oit1*. **e,** CloserRegions (Regions 34 and 35) for *Fhit*.

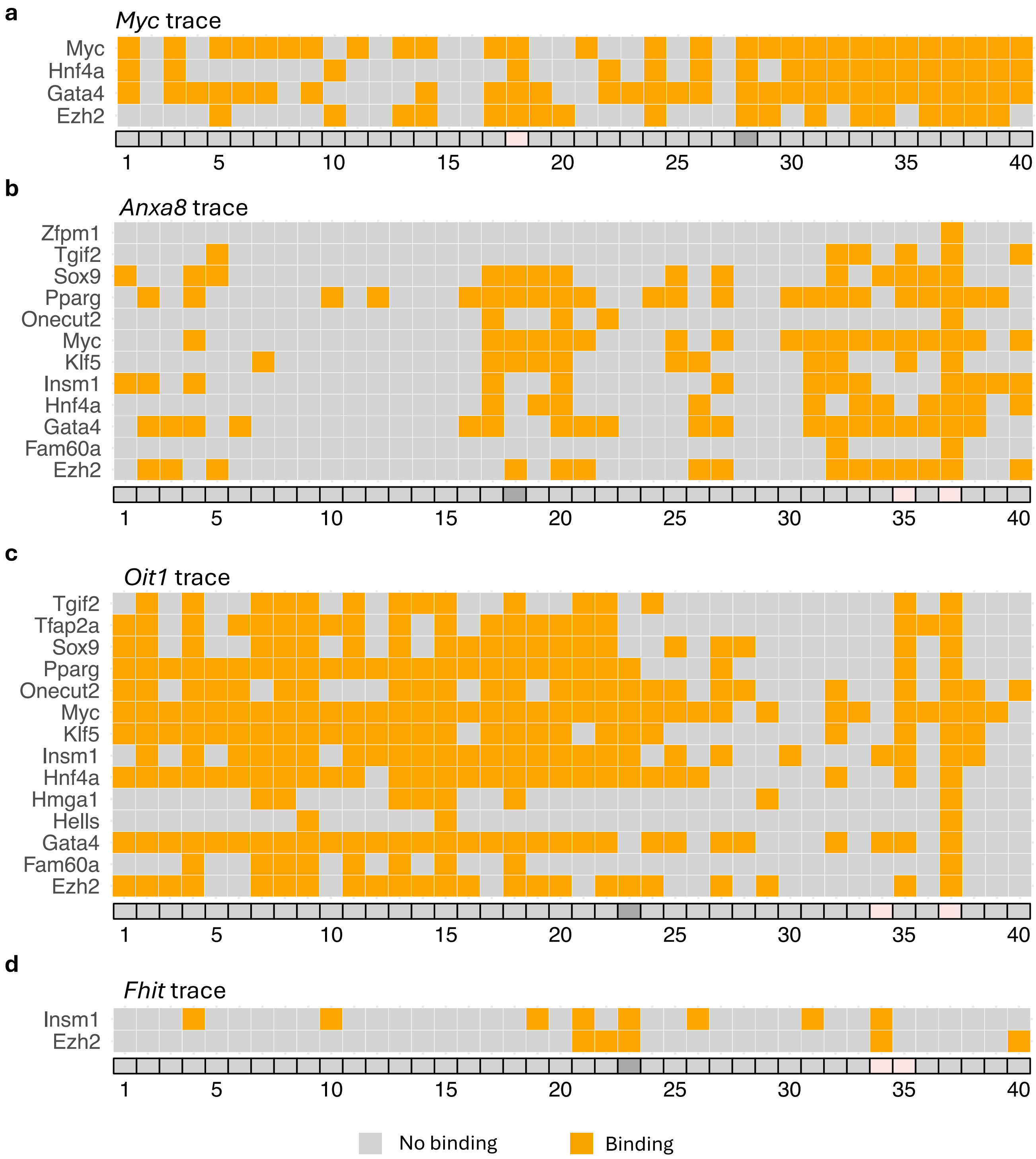


**Supplementary Fig. 10. Regulator (y-axis) binding sites on CloserRegions (pink regions on x-axis) for each CPD trace. a,** for *Myc*. **b,** for *Anxa8*. **c,** for *Oit1*. **d,** for *Fhit.* Orange boxes indicate binding; gray boxes indicate no binding.

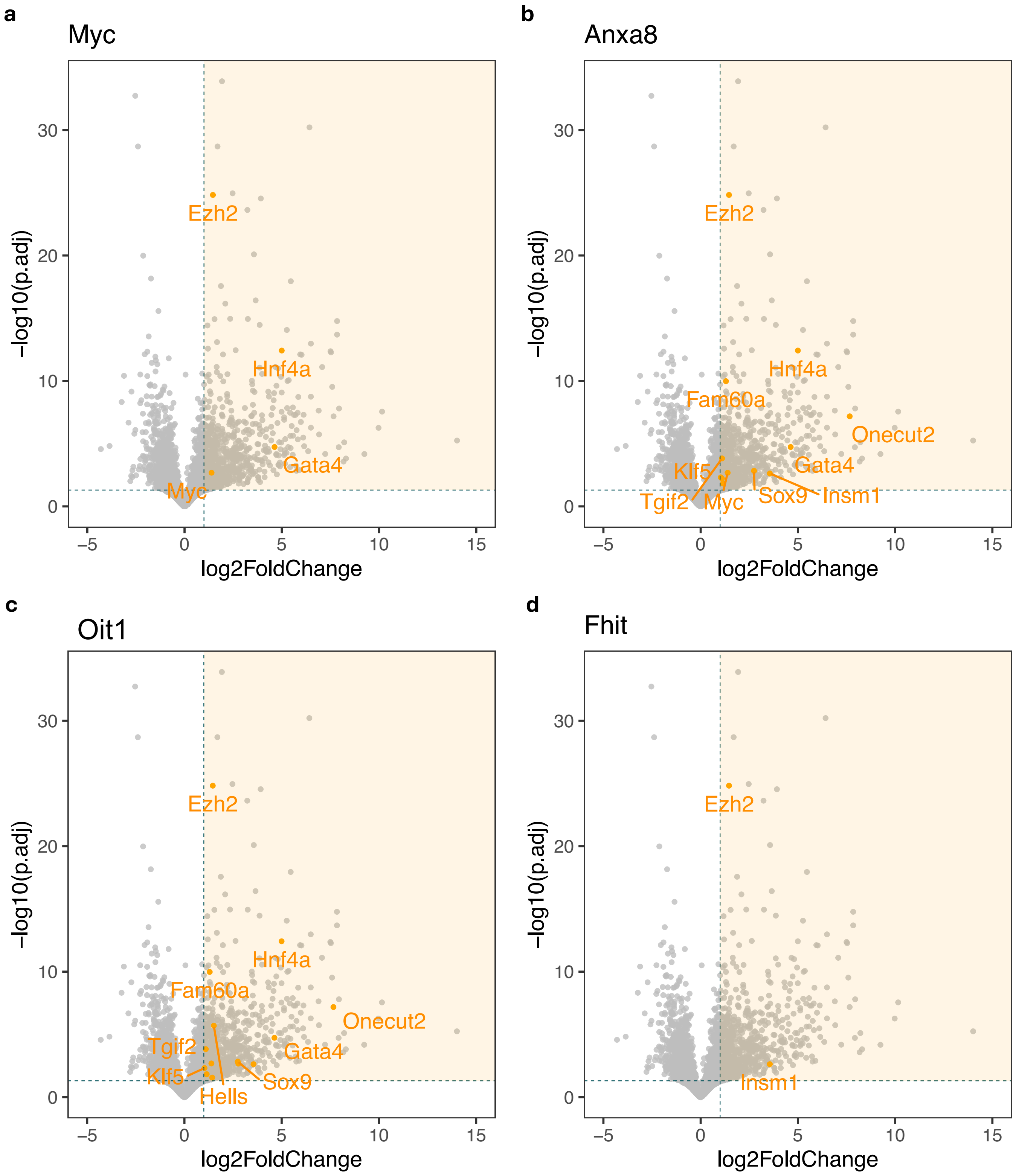


**Supplementary Fig. 11. Volcano plot showing the differential gene expression between LUAD and adenoma.** The shaded region highlights genes significantly upregulated in LUAD than in adenoma (log2FoldChange > 1 & adj.p < 0.05). Orange dots indicate the upregulated TR genes that potentially bind to the CloserRegions in the CPD trace. **a,** for *Myc*. **b,** for *Anxa8*. **c,** for *Oit1*. **d,** for *Fhit*.

**a**

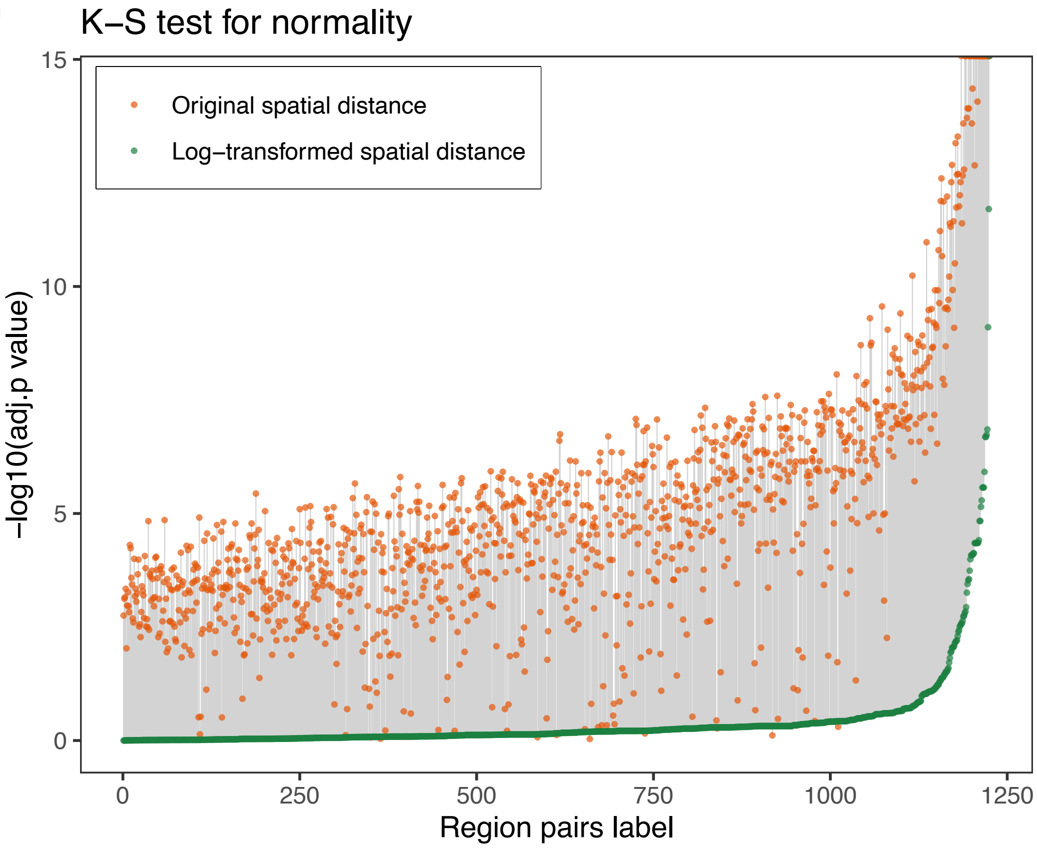


**b**

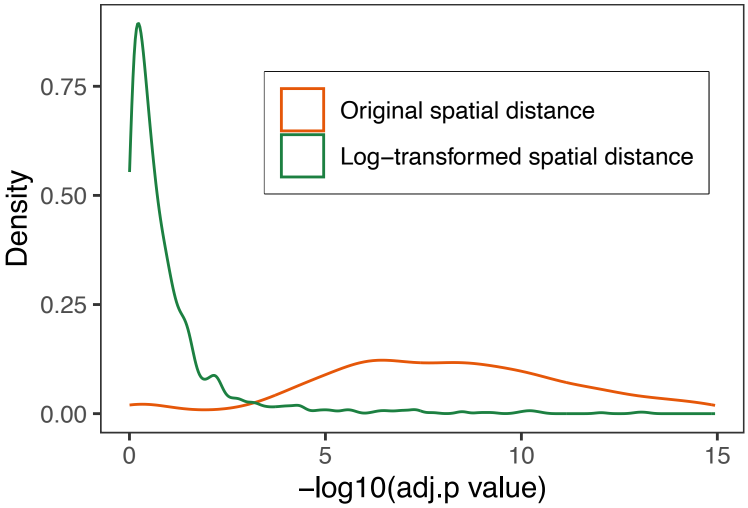


**Supplementary Fig. 12. Assessment of chromatin tracing data normality for the original data and the logarithmic transformed data using K-S test.** The null hypothesis of K-S test here is that the sample follows the normal distribution. **a,** -log10(adj.p) of K-S test for each region pair's original spatial distance (orange dots) and log-transformed spatial distance (green dots). The gray line connecting the orange dot and the green dot to indicate the p-value change after log-transforming of the data**. b,** the density distribution of –log10(adj.p) for using the original spatial distance (orange line) and the log-transformed spatial distance (green line).
